# Mapping subcortical fear pathways in the human brain: thalamo-amygdala connections revealed by high-resolution tractography

**DOI:** 10.64898/2026.01.07.697914

**Authors:** Emmanouela Kosteletou-Kassotaki, Liu Mengxing, Martina T. Cinca-Tomás, Pedro M. Paz-Alonso, Judith Domínguez-Borràs

## Abstract

Influential models of emotion have postulated the existence of several direct subcortical pathways, or ‘low roads’, that convey fast sensory thalamic inputs to the amygdala for affective processing. In rodents and non-human primates, specific thalamo-amygdala connections have been identified and linked to fear responses. These connections mainly originate in three thalamic nuclear groups (posterior, intralaminar, and medial) and project to the basolateral amygdala (BLA). However, the existence of similar thalamo-amygdala pathways in humans remains uncertain. Here, aiming to reproduce subcortical amygdala connections previously described in the non-human animal literature, we mapped those tracts in 113 human participants of either sex. We implemented an advanced high-resolution tractography protocol optimized for reliably tracing axonal pathways through regions with complex white-matter architecture. We reconstructed white matter tracts connecting thalamic nuclei to the BLA, and assessed their test-retest reproducibility to evaluate their anatomical plausibility. Among posterior thalamic nuclei, projections from the medial geniculate nucleus and the medial and inferior pulvinar to the BLA emerged as the strongest and most robust thalamo-amygdala pathways. In turn, the mediodorsal nucleus within the medial group exhibited a prominent connection to the BLA, with a high number of streamlines and moderate-to-high reliability across sessions. Finally, intralaminar thalamic nuclei (parafascicular, central medial, and centromedian) showed consistent connections to the BLA, albeit with higher intrasubject variability and moderate-to-low reproducibility. Together, these findings provide a unifying anatomical framework for multiple direct thalamo-amygdala pathways in the human brain, and suggest the existence of distinct evolutionarily conserved routes for subcortical affective processing.

**Significance statement:** Animal studies have identified several subcortical “low road” pathways that may convey direct sensory inputs from the thalamus to the amygdala. Here, we analyzed human diffusion magnetic resonance imaging data to reconstruct thalamo-amygdala pathways previously reported in non-human species and assessed their test-retest reproducibility. White matter tracts linking the medial geniculate nucleus, as well as the medial and inferior pulvinar, to the basolateral amygdala appeared to be the most robust and reliable. Additional connections were found between the mediodorsal and intralaminar nuclei of the thalamus and the basolateral amygdala. These findings reveal the presence of multiple conserved direct pathways to the amygdala in humans, likely supporting rapid affective processing across modalities.

## Introduction

Rapid processing of emotional signals is a key evolutionary adaptation for vertebrates. Previous work in rodents, cats, and non-human primates has identified several subcortical neural connections, or ‘low roads’, which directly connect the thalamus and the amygdala, presumably associated with rapid fear processing and adaptive behavior (Carr, 2015; McFadyen et al., 2020). Specifically, the basolateral amygdala (BLA), a key structure for fear conditioning and other affective processes, has been described as a major target of thalamic projections associated with affective function (see Janak & Tye, 2015; Pessoa & Adolphs, 2010; Phelps & LeDoux, 2005; Wassum & Izquierdo, 2015). Converging evidence from non-human animals suggests that the BLA receives sensory afferents from three main thalamic subregions, which include posterior, intralaminar, and midline nuclei, via projections implicated in fear-related processes (see Table 1).

**Table 1:**
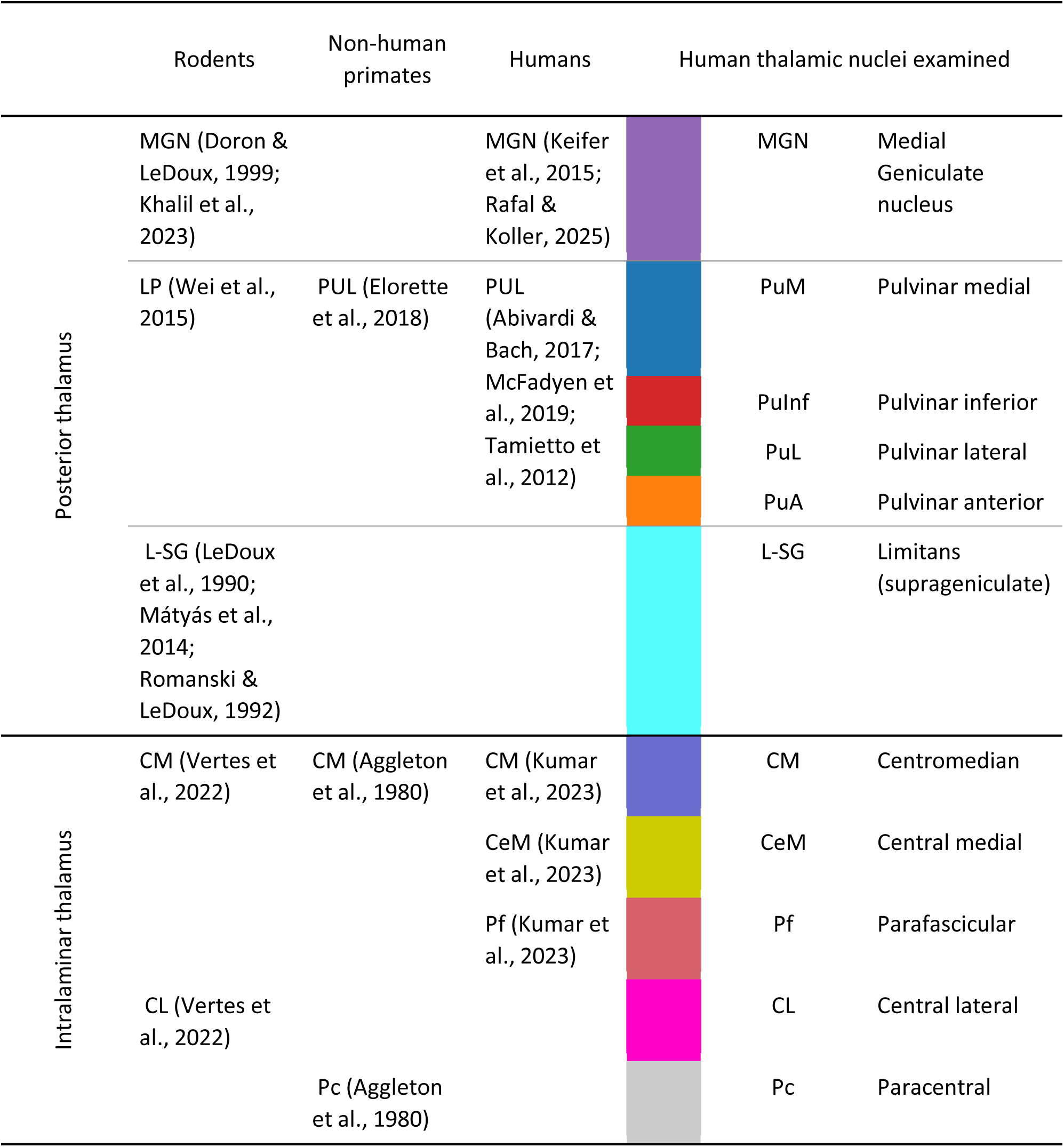
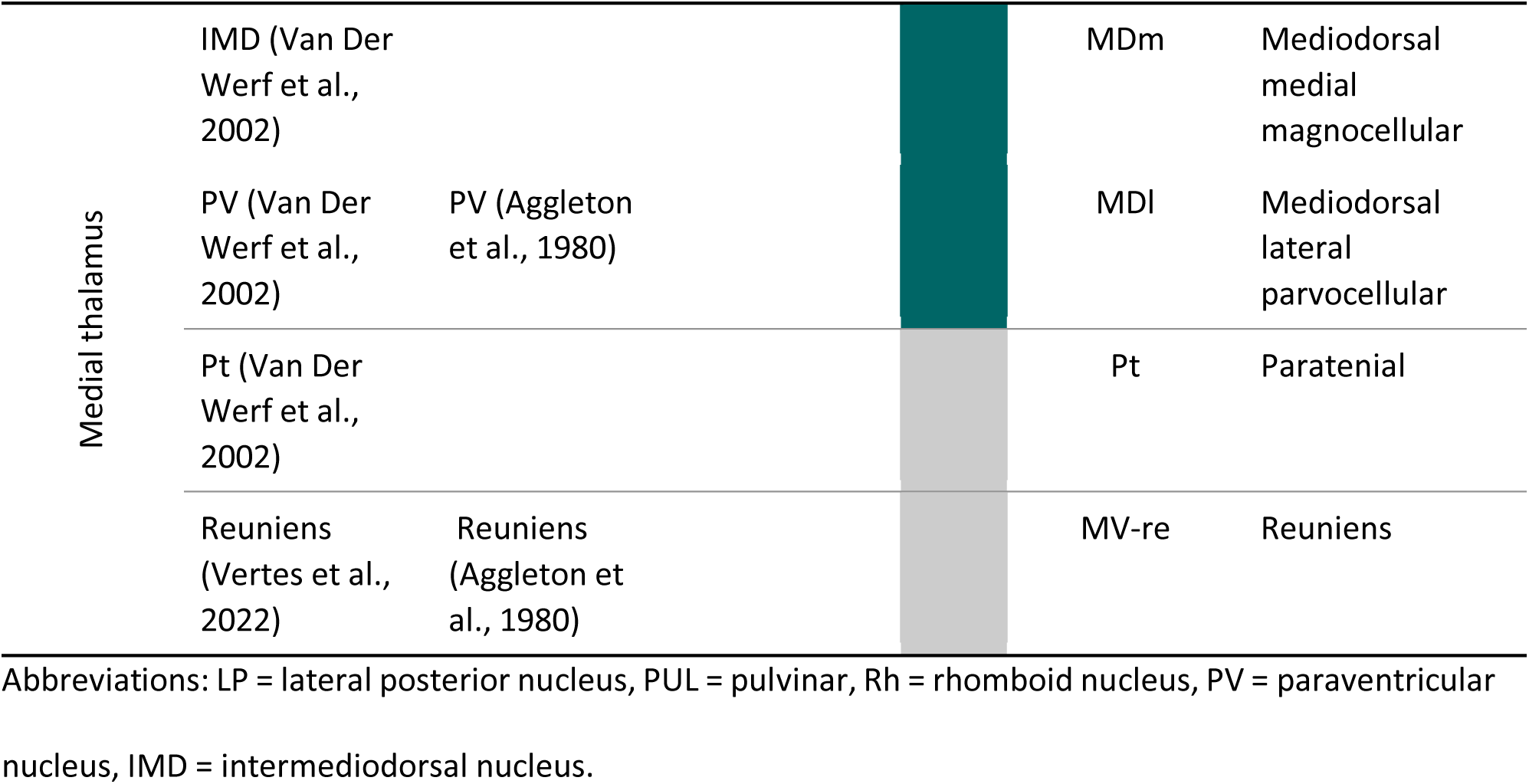
Summary of thalamic nuclei reported in the literature to project to the basolateral amygdala across species (rodents, non-human primates, and humans), including the putative human homologues of the regions identified in animal studies. The last three columns indicate the specific thalamic nuclei included in our analysis, with colors corresponding to those shown in Figure 1A.

Within the posterior thalamus, fear-related pathways include projections from the medial geniculate nucleus (MGN), a major auditory relay (Doron & LeDoux, 1999; LeDoux et al., 1984; Shinonaga et al., 1994), from the suprageniculate nucleus, a non-lemniscal thalamic region involved in the integration of sensory and affective signals (LeDoux et al., 1990; Romanski & LeDoux, 1992), and from the primate pulvinar (i.e. the lateral posterior nucleus in rodents and cats), a key region for visual attention and multisensory processes (Day-Brown et al., 2010; Elorette et al., 2018; Grieve et al., 2000; Wei et al., 2015). In humans, while MGN- and suprageniculate-amygdala connectivity remains poorly characterized, a pulvinar-BLA pathway has been extensively reported and implicated in fear-related processing (Celeghin et al., 2019; Domínguez-Borràs & Vuilleumier, 2022; Kragel et al., 2021; McFadyen et al., 2019; Tamietto et al., 2012; Tamietto & De Gelder, 2010). However, the pulvinar is a large thalamic complex comprising multiple functionally specialized subdivisions (Bourgeois et al., 2020; Froesel et al., 2024) and their specific anatomical connectivity with human amygdala remains undefined. Finally, intralaminar and midline thalamic projections to the BLA have been consistently reported in rodents, cats and non-human primates, and associated with fear-related and defensive behaviors (Aggleton et al., 1980; Dobi et al., 2013; Mátyás et al., 2014; Ottersen & Ben-Ari, 1979; Su & Bentivoglio, 1990), but again are largely unknown in humans. Taken together, evidence across species suggests that multiple evolutionarily conserved thalamo-amygdala pathways for fear may exist in humans (Carr, 2015; McFadyen et al., 2020), but their anatomical specificity and organizational structure remain uncharacterized.

Here, we sought to systematically map thalamo-amygdala pathways in the human brain that have been previously described and implicated in fear processes in non-human species. To this end, we reconstructed white matter tracts connecting key thalamic nuclei to the basolateral amygdaloid complex using diffusion magnetic resonance imaging (MRI) in a sample of 113 participants. Mapping connectivity around these deep subcortical structures in the human brain is restricted by the dense white-matter architecture and complex fiber crossings around them, which can limit anatomical specificity in the tract reconstruction (Jeurissen et al., 2014). To mitigate this constraint, we used an advanced tractography protocol designed to maximize reliability and validity of the reconstructed tracts. This protocol has already been used to reliably map human first-order-relay thalamic connectivity (Liu et al., 2022) and mediodorsal thalamus to prefrontal cortex pathways (Mengxing et al., 2023), by leveraging tools capable of discerning crossing fibers. Moreover, to provide a spatially detailed anatomical overview of the tracts, we used probabilistic thalamic (Iglesias et al., 2018) and high-resolution amygdala (Saygin et al., 2017) parcellation atlases. Finally, to further assess the reliability and, by extension, the anatomical plausibility of the reconstructed pathways, we compared their white matter microstructural and macrostructural properties across test-retest sessions in a subset of participants who underwent a second MRI session. Our results outline previously uncharted subcortical pathways potentially relevant to human emotion.

## Methods

### Subjects

A total of 113 healthy, right-handed volunteers (65 females; mean age = 24.5 years, SD = 4.33; 65 females) participated. All had normal or corrected-to-normal vision and no history of major medical, neurological, or psychiatric disorders. A subset of 24 participants (mean age = 24.7 years, SD = 4.06; 13 females) returned for a second session, undergoing the same MRI protocol after a mean interval of 15 days (SD = 21.82, range: 7-104 days). The study was approved by the Ethics Committee of the Basque Center on Cognition, Brain and Language (BCBL) and conducted in accordance with the Code of Ethics of the World Medical Association (Declaration of Helsinki). Before participation, all individuals provided written informed consent and received monetary compensation for their involvement.

### Data acquisition

Whole-brain MRI data were acquired using a 3T Siemens Prisma Fit scanner (Siemens Medical Solutions) equipped with a 64-channel whole-head coil. The MRI protocol included a high-resolution T1-weighted (T1w) MPRAGE structural scan and DWI sequences (described in Liu et al., 2022). The T1-weighted structural images were obtained with the following parameters: repetition time (TR) = 2530 ms, echo time (TE) = 2.36 ms, flip angle (FA) = 7°, field of view (FOV) = 256 mm, voxel size = 1 mm isotropic, and 176 slices. For diffusion imaging, a total of 100 diffusion-weighted images were collected in the anterior-to-posterior phase-encoding direction, including 50 images at a b-value of 1000 s/mm² and another 50 at a b-value of 2000 s/mm². Additionally, 12 non-diffusion-weighted (b = 0 s/mm²) images were acquired for motion and geometrical distortion correction (five with the same phase-encoding direction as the DWI images and seven with the reversed phase-encoding direction; posterior-to-anterior). Both the diffusion-weighted and b0 images were acquired using the following parameters: TR = 3600 ms, TE = 73 ms, FA = 78°, voxel size = 2 mm isotropic, 72 slices without gap, and a multiband acceleration factor of 3 (see Liu et al., 2022).

### Tractography pipeline

Tract reconstruction was performed using the Reproducible Tract Profiles 2 (RTP2) pipeline (Lerma-Usabiaga et al., 2023), ensuring data provenance and reproducibility. The RTP2 pipeline organizes the process into region of interest (ROI) definition, DWI preprocessing and tract identification and tractometry. A total of 10 white matter tracts were reconstructed bilaterally, connecting different thalamic subregions with the lateral, basal, and accessory basal nuclei of the amygdala.

### ROI definition

The first step of the RTP2-pipeline (RTP2-anatROIs) involves processing each subject’s T1w anatomical image to obtain the ROIs for tractography. This process takes as input the T1w image of the subject and predefined ROIs in MNI space, and outputs a segmented T1w image along with the ROIs in individual subject T1w space.

First, Freesurfer (http://surfer.nmr.mgh.harvard.edu/) was used to perform cortical-subcortical segmentation of the T1w image and define the thalamic and amygdala nuclei. To obtain the thalamic nuclei, we applied FreeSurfer’s thalamic segmentation module, which leverages a probabilistic atlas built from histological and high-resolution *ex vivo* MRI data (Iglesias et al., 2018). The thalamic nuclei selected as ROIs in this study included the medial geniculate nucleus (MGN), the pulvinar (PUL), the suprageniculate nucleus (L-SG) and the entire intralaminar and medial nuclei. It is important to note that the thalamic segmentation atlas provided by Iglesias et al., (2018) divides the PUL into four subregions: the medial (PuM), inferior (PuInf), lateral (PuL) and anterior (PuA) pulvinar. Similarly, the intralaminar nucleus is divided into five subregions: the central medial (CeM), centromedian (CM), central lateral (CL), parafascicular (Pf) and paracentral (Pc) and the medial nucleus into four subregions: the mediodorsal medial (MDm), mediodorsal lateral (MDl), paratenial (Pt), and reuniens (MV-re). While rodent atlases distinctly segregate mediodorsal (MD), paratenial, and reuniens nuclei into separate medial and midline groups, the human atlas we employed (Iglesias et al., 2018) classifies them collectively within a broader medial thalamic group. In our analysis, we combined the mediodorsal medial (MDm) and mediodorsal lateral (MDl) nuclei into a single ROI, and all subsequent analyses were conducted on this unified subregion for better reproducibility. For the BLA nuclei, we used the amygdala segmentation by Saygin et al., (2017), implemented in FreeSurfer, including its lateral, basal, and accessory basal subregions. Lastly, the inferior colliculus (IC) ROI was sourced from the Brainstem Navigator toolkit (Bianciardi et al., 2023), which provides in-vivo atlas labels for brainstem nuclei based on semi-automatic and manual segmentations of high-resolution, multi-contrast 7 Tesla MRI data. The IC ROI was transformed from MNI space to individual space using a nonlinear transformation in Advanced Normalization Tools (ANTs; http://stnava.github.io/ANTs). To ensure that the ROIs extended to the gray-white matter interface, we dilated them by one voxel. Considering the anatomical location and expected connectivity of the tracts of interest, we employed inclusion and exclusion ROIs to enhance neuroanatomical accuracy during tract reconstruction, following an initial visual inspection. Table 1 summarizes thalamic nuclei reported to project to the BLA across species, based on histological tract-tracing, optogenetic, and neuroimaging (diffusion MRI tractography) evidence from rodents, non-human primates, and humans. Putative human homologues of the thalamic regions identified in animal studies are also indicated. Figures 1A, B show the thalamic and amygdala ROIs used in our study.

**Figure 1:**
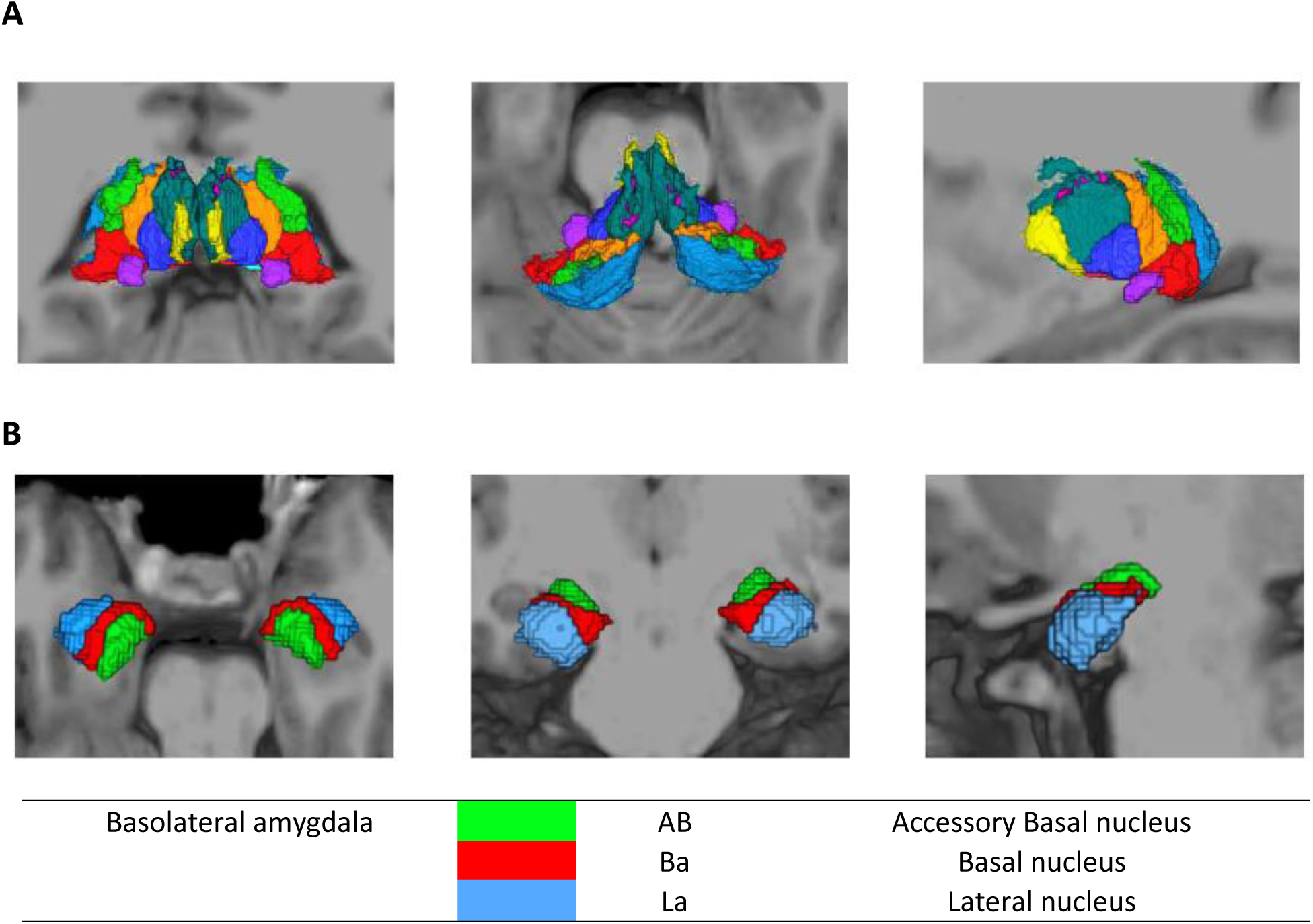
Thalamic and amygdala parcellations used in our study. (**A**) Thalamic parcellations. Coronal, axial and sagittal slices (left to right, respectively) of the thalamic nuclei, as produced by the Iglesias et al., (2018) atlas in a representative subject. Colors denote thalamic nuclei listed in Table 1; grey indicates nuclei that were too small to be visually distinguishable. (**B**) Amygdala parcellations defining the basolateral amygdala. Coronal, axial and sagittal slices (left to right, respectively) of the accessory basal (green), basal (red) and lateral (light blue) nuclei of the amygdala, as produced by the Saygin et al., (2017) atlas in a representative subject.

### DWI data preprocessing

After obtaining the ROIs that were used in the study, the second step of the RTP2 pipeline (RTP2-preproc) was implemented to preprocess diffusion data and registering it in anatomical space. This step followed recommendations from MRtrix3 (Tournier et al., 2019) and utilized tools from MRtrix3, ANTs, and FSL (Jenkinson et al., 2012). The preprocessing was carried out through several steps using MRtrix3 functions. First, data denoising was performed based on random matrix theory, leveraging data redundancy in the patch-level principal component analysis domain (Cordero-Grande et al., 2019; Veraart et al., 2016) using the dwidenoise function. Second, Gibbs ringing artifacts were corrected using mrdegibbs (Kellner et al., 2016). Third, susceptibility-induced distortions and motion artifacts were corrected using FSL’s topup and eddy tools (Smith et al., 2004), called by dwifslpreproc. Fourth, B1 field inhomogeneity correction was performed using dwibiascorrect, and Rician background noise was removed with mrcalc. Finally, a rigid transformation matrix was applied to align the DWI images to the corresponding T1w using ANTs. To ensure that the diffusion data from test and retest sessions were in the same space, images from both sessions were aligned to the same T1w scan collected during the initial test session. This allowed both test and retest sessions to use the same ROIs, while only diffusion preprocessing and streamline tracking remained session-specific.

### Tract identification and tractometry

In the third and final step, the RTP2-pipeline was used to generate the final white matter tracts. This step used the ROIs and preprocessed DWI data to systematically identify the tracts of interest bilaterally. To assess the robustness and anatomical plausibility of the tracts, we first reconstructed the pathways between all thalamic ROIs described above, and the BLA in a randomly selected subset of four subjects. After visually inspecting each tract, we refined the iteration parameters, including the placement of additional exclusion ROIs, if necessary, to ensure that reconstructed tracts did not overlap with those from neighboring thalamic nuclei projecting to the amygdala. Following these adjustments and after visually inspecting the generated tracts, we excluded those that did not appear anatomically reliable. Additionally, we investigated the possibility of a direct pathway connecting the IC with BLA, bypassing the MGN, based on tract-tracing evidence of this connection in bats (Marsh et al., 2002). The final reconstructed pathways and the specific ROIs used are detailed in Table 2. For all tracts, streamlines were first seeded from inclusion ROI 1 and terminated in inclusion ROI 2. Then, a second batch of streamlines was generated by seeding from ROI2 and terminating in ROI 1. The final tract was obtained by combining streamlined from both directions.

**Table 2:**
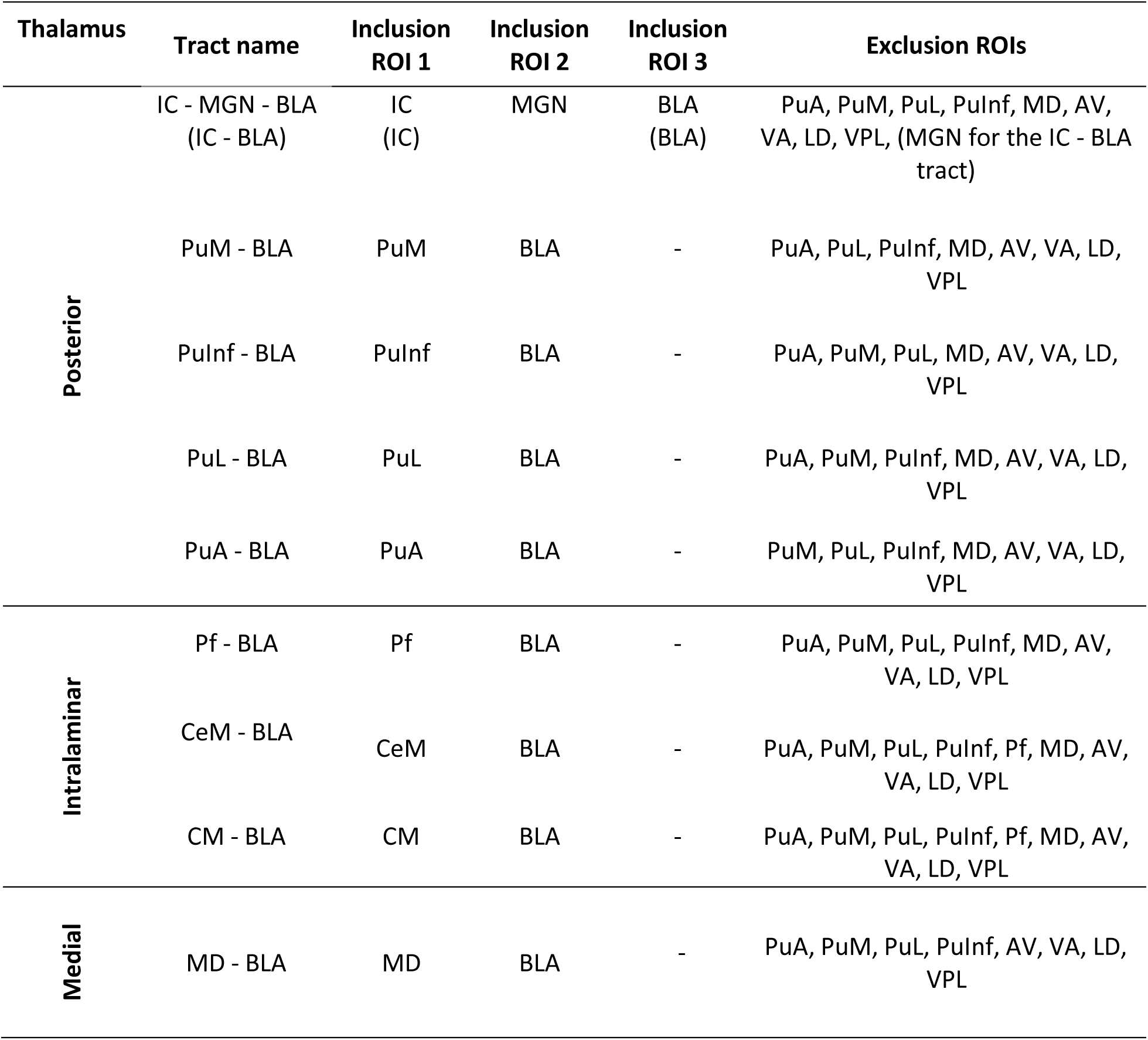
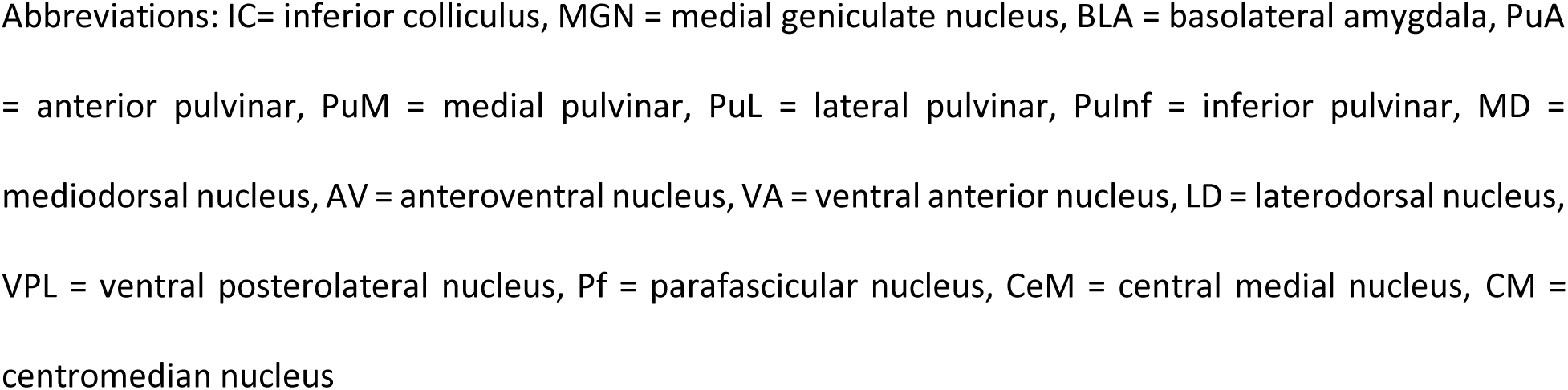
Tracts and ROI constraints for tract reconstruction.

| Thalamus | Tract name | Inclusion ROI 1 | Inclusion ROI 2 | Inclusion ROI 3 | Exclusion ROIs |
| --- | --- | --- | --- | --- | --- |
| Posterior | IC - MGN - BLA<br>(IC - BLA) | IC<br>(IC) | MGN | BLA<br>(BLA) | PuA, PuM, PuL, PuInf, MD, AV, VA, LD, VPL, (MGN for the IC - BLA tract) |
|  | PuM - BLA | PuM | BLA | - | PuA, PuL, PuInf, MD, AV, VA, LD, VPL |
|  | PuInf - BLA | PuInf | BLA | - | PuA, PuM, PuL, MD, AV, VA, LD, VPL |
|  | PuL - BLA | PuL | BLA | - | PuA, PuM, PuInf, MD, AV, VA, LD, VPL |
|  | PuA - BLA | PuA | BLA | - | PuM, PuL, PuInf, MD, AV, VA, LD, VPL |
| Intralaminar | Pf - BLA | Pf | BLA | - | PuA, PuM, PuL, PuInf, MD, AV, VA, LD, VPL |
|  | CeM - BLA | CeM | BLA | - | PuA, PuM, PuL, PuInf, Pf, MD, AV, VA, LD, VPL |
|  | CM - BLA | CM | BLA | - | PuA, PuM, PuL, PuInf, Pf, MD, AV, VA, LD, VPL |
| Medial | MD - BLA | MD | BLA | - | PuA, PuM, PuL, PuInf, AV, VA, LD, VPL |

Abbreviations: IC= inferior colliculus, MGN = medial geniculate nucleus, BLA = basolateral amygdala, PuA = anterior pulvinar, PuM = medial pulvinar, PuL = lateral pulvinar, PuInf = inferior pulvinar, MD = mediodorsal nucleus, AV = anteroventral nucleus, VA = ventral anterior nucleus, LD = laterodorsal nucleus, VPL = ventral posterolateral nucleus, Pf = parafascicular nucleus, CeM = central medial nucleus, CM = centromedian nucleus

For the IC-MGN-BLA tract, apart from the seed and terminal ROIs IC and BLA, a third inclusion MGN ROI was included to ensure that all streamlines passed through the MGN. This constraint was applied because this corresponds to the subcortical pathway originally described in animal models (Doron & LeDoux, 1999; LeDoux et al., 1984), where auditory information is relayed from IC to MGB and subsequently to BLA. It also allows for comparison with a direct IC-BLA tract, in which we used MGN as an exclusion ROI. Exclusion ROIs were used to ensure that streamlines passing through these regions were not included in the tract reconstruction, so as to avoid potential confounds in the identification of pathways.

We first modeled the diffusion data to generate fiber orientation distributions (FODs), which represent possible fiber directions with corresponding weights in each voxel. This was done using MRtrix3’s multi-tissue constrained spherical deconvolution (CSD; Jeurissen et al., 2014), allowing for the identification of crossing fibers and the estimation of multiple directions per voxel. Next, streamline tractography was performed on the estimated FODs using the probabilistic iFOD2 algorithm (Tournier et al., 2012) with the following parameters: step size 1 mm, maximum fiber length 80 mm, minimum fiber length 10 mm, FOD amplitude threshold of 0.1, and angle threshold of 45 degrees. We generated along-tract profiles for fractional anisotropy (FA) using the tract metrics obtained from Vistasoft. To do this, we first identified the central location of all streamlines within each tract and divided it into 100 equal-length segments. FA values were then sampled within a disc centered on each segment, and a weighted average was calculated to summarize the diffusion properties of that region.

### Statistical analysis and reproducibility measurements

We ran the protocol on the dataset of 113 participants reconstructing the 9 thalamus-basolateral amygdala tracts and the additional direct IC-BLA tract. To examine tract-specific and hemispheric differences in the number of streamlines, two-way repeated measures ANOVAs were conducted separately for each thalamic subgroup (posterior, intralaminar), with pathway and hemisphere as within-subject factors, and the number of streamlines as the dependent variable. Subject was included as a random factor to account for within-subject variability. The assumption of sphericity was assessed using Mauchly’s test, and when violated, Greenhouse-Geisser corrections were applied to adjust the degrees of freedom. To further explore significant interaction effects, post hoc pairwise comparisons were conducted using the emmeans package (Lenth R., 2025). Specifically, pairwise comparisons between the left and right hemispheres were performed within each tract using estimated marginal means, and Bonferroni correction was applied to adjust for multiple comparisons. For both the medial subgroup, which included only the MD-BLA tract, and the IC-BLA tract, two separate paired sample t-tests were performed to compare hemispheric differences in streamline counts.

To examine the reliability of the reconstructed tracts we tested their reproducibility across two scanning sessions, on a subset of 24 participants, who returned for a second acquisition session within a mean temporal interval of 15 days. We examined how consistently our protocol reconstructed the tracts and tractometry at these two different points in time. To measure the test-retest reproducibility for the two MRI sessions, we quantitatively analyzed the similarity of two tracts at both microstructural and macrostructural levels. At the microstructural level, to evaluate reproducibility in tractometry generation, we performed pairwise Pearson correlations on tract profiles across test and retest sessions. We report the correlations for fractional anisotropy (FA) values, as this is the most widely used metric in the literature. At the macrostructural level, we calculated: 1) the Dice overlap index to evaluate the volume-based overlap between test and retest sessions, 2) the bundle adjacency to calculate the average distance between streamlines from two tracts, and 3) the streamline density correlation for the voxel-level streamline density of all tract pairs. Together, these metrics provide a comprehensive assessment of macrostructural reproducibility, as each capture distinct properties of tract organization. However, it is important to note that these indices are inherently sensitive to the number of reconstructed streamlines. When streamline counts are low, small spatial deviations can disproportionately reduce Dice overlap and density correlations, and bundle adjacency estimates may become unstable. Consequently, lower reproducibility values for sparsely reconstructed tracts should be interpreted with caution, as they may reflect methodological constraints related to tract sparsity rather than true anatomical variability. This analysis was performed using the package scilpy (see details in Schilling et al., (2021) and https://github.com/scilus/scilpy), and was conducted for each tract.

## Results

### White matter tracts connecting posterior, intralaminar and medial thalamic subregions to the basolateral amygdala

Our goal was to map the anatomical connections between the thalamus and the BLA in humans, based on converging evidence from non-human animals. We used a high-resolution tractography protocol, optimized for a reproducible mapping of thalamic pathways (Lerma-Usabiaga et al., 2023; Liu et al., 2022; Mengxing et al., 2023), to reconstruct BLA projections from three major thalamic subregions, comprising the posterior, the intralaminar and the medial nuclei of the thalamus. To further assess the micro- and macrostructural reliability of the reconstructed pathways, we employed a test-retest reproducibility approach. DWI data were acquired from the same 24 participants from our total sample across two separate sessions, using identical MRI protocols. Overall, we defined connections as robust if they were detected in the majority of participants with a high number of streamlines and high reproducibility, and as weak when the number of streamlines was low and reproducibility was moderate or low.

#### Robust connections between the posterior thalamus and the BLA

Posterior thalamic tracts that survived initial inspection included those originating from the MGN and the pulvinar (PUL). To improve anatomical precision for the MGN tract (Bordi & LeDoux, 1994), we reconstructed the white matter pathway linking IC to MGN and subsequently to BLA. In addition, we reconstructed distinct tracts from each of the four pulvinar subregions - medial (PuM), inferior (PuInf), lateral (PuL) and anterior (PuA) - to the BLA.

All 113 participants exhibited at least one streamline connecting the PuM-BLA, PuInf-BLA and PuL-BLA pathways. Conversely, the IC-MGN-BLA pathway was successfully reconstructed in 96.5% (left hemisphere) and 92.9% (right hemisphere) of participants, whereas the PuA-BLA tract was identified in 78.8% (left) and 77.0% (right). Based on both the proportion of successful reconstructions and streamline counts, the IC-MGN-BLA, PuInf-BLA, PuM-BLA, PuL-BLA pathways were considered robust, whereas the PuA-BLA pathway was considered weak. ANOVA results showed a significant main effect of pathway (F (3.09, 346.05) = 5382.99, p < 0.001, η^2^ = 0.95), and hemisphere (F (1, 112) = 12.56, p < 0.001, η^2^ = 0.012), as well as a significant pathway x hemisphere interaction (F (2.61, 292.4) = 4.87, p = 0.004, η^2^ = 0.014), indicating that hemispheric differences in the number of streamlines differed across pathways (see Table 3). Post-hoc pairwise comparisons between hemispheres within each tract (Bonferroni corrected) revealed a significant hemispheric lateralization in the IC-MGN-BLA and PuInf-BLA pathways, with more streamlines in the left compared to the right hemisphere (b = 252.05, SE = 58.7, t _(112)_ = 4.3, p < 0.001; b = 240.63, SE = 59.8, t _(112)_ = 4.03, p = 0.0001, respectively). No significant hemispheric differences were found for the PuM-BLA, PuL-BLA and PuA-BLA pathways (all p > 0.05).

**Table 3:**
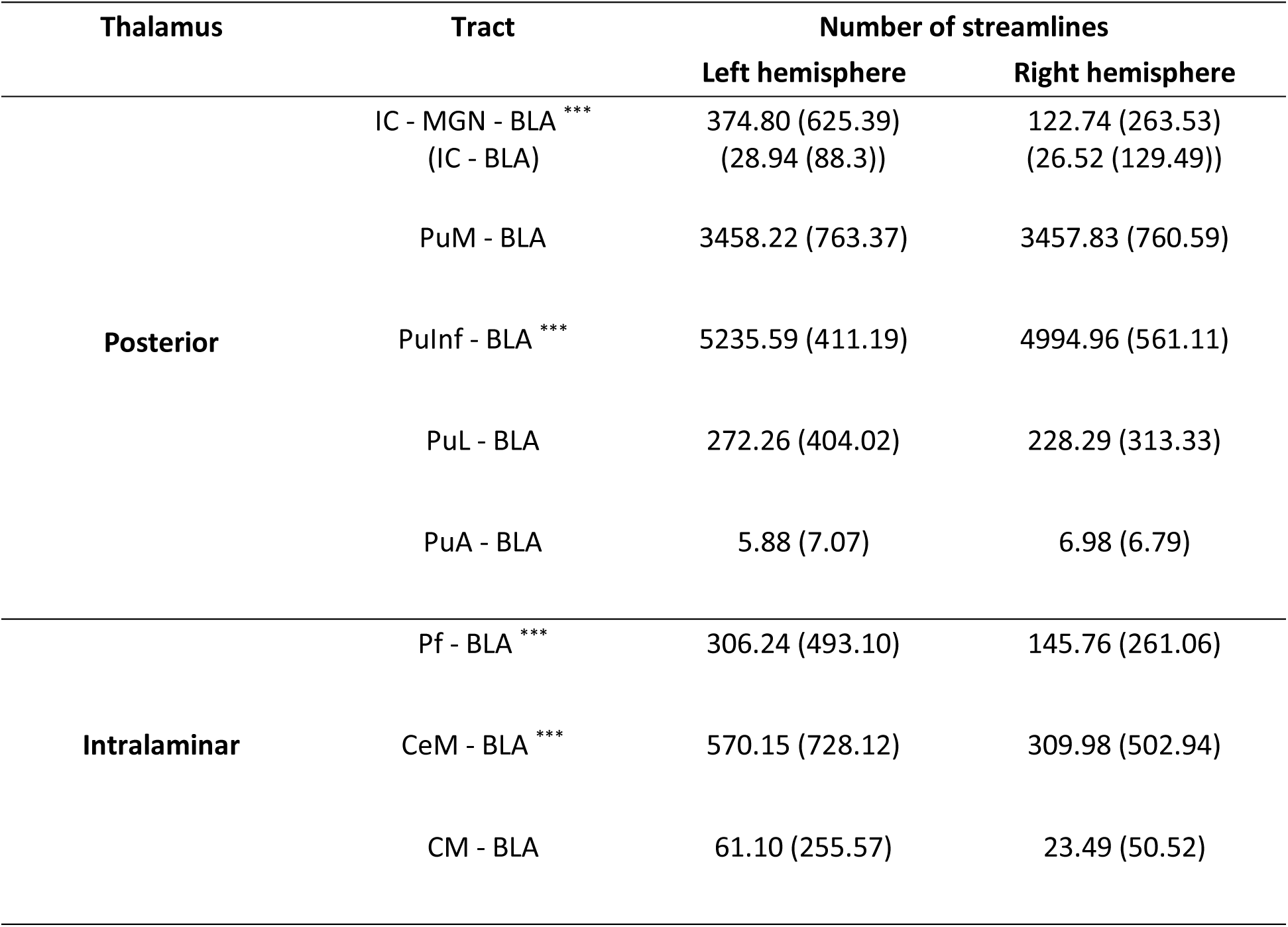
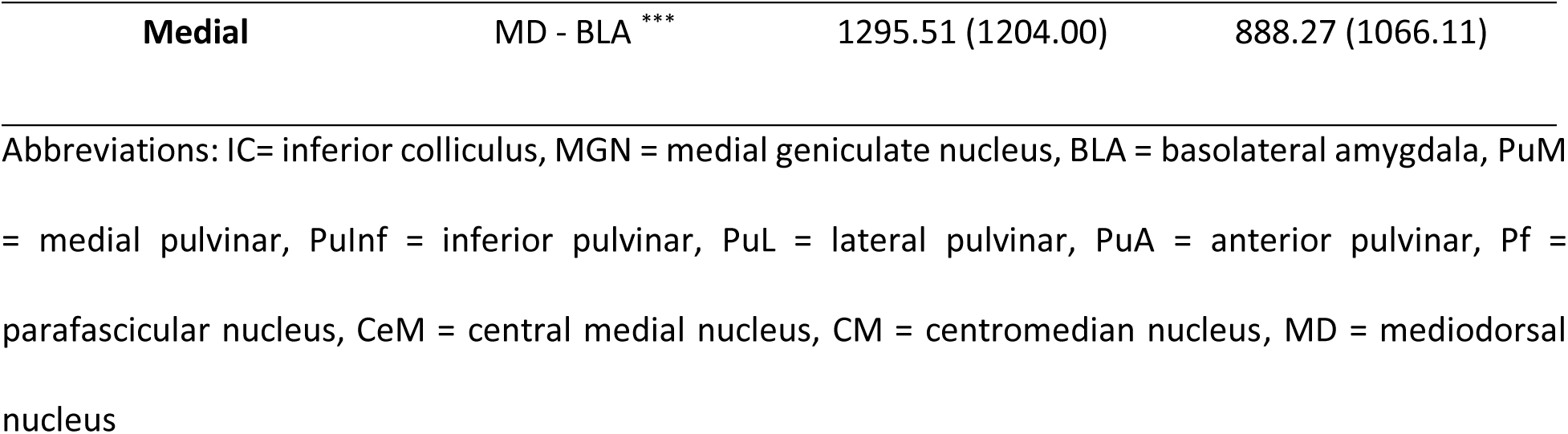
Streamline counts (± SD) for the posterior, intralaminar and medial thalamic subdivisions. Pathways with significant hemispheric lateralization are marked with asterisk: *** p < 0.001.

Notably, PuM and PuInf exhibited robust bilateral connections to the BLA, whereas PuL showed fewer streamlines. In contrast, PuA, when present, showed only a sparse number of connecting streamlines.

#### Consistent but variable connections between the intralaminar thalamus and BLA

After visually inspecting the intralaminar-BLA tracts previously described in the literature (see Table 1), in a small subset of participants, we excluded the pathways that appeared structurally unreliable, including central lateral (CL) and paracentral (Pc). The parafascicular (Pf), central medial (CeM), and centromedian (CM) nuclei demonstrated consistent tract reconstructions, allowing us to generate streamline connections between these nuclei and the BLA across the entire cohort. The number of streamlines varied across participants. While most individuals exhibited measurable connections, specific pathways were not detected in some cases. Specifically, the Pf-BLA tract was reconstructed successfully in 97.3% and 96.5% of the participants in the left and right hemispheres, respectively; the CeM-BLA in 98.2% and 96.5; and the CM-BLA in 88.5% and 84.1%. Based on both reconstruction prevalence and streamline counts, the Pf-BLA and CeM-BLA pathways were considered relatively robust, whereas the CM-BLA pathway appeared less robust. ANOVA results on the intralaminar group revealed significant main effects of pathway (F (1.48, 165.3) = 61.14, p < 0.001, η^2^ = 0.12), and hemisphere (F (1, 112) = 18.74, p < 0.001, η^2^ = 0.03), as well as a significant pathway x hemisphere interaction (F (1.81, 202.7) = 9.87, p < 0.001, η^2^ = 0.011; see Table 3). Post hoc pairwise comparisons revealed significant hemispheric differences in the Pf-BLA and CeM-BLA pathways, with greater number of streamlines in the left hemisphere compared to the right (b = 160.5, SE = 46, t _(112)_ = 3.49, p = 0.0007; b = 260.2, SE = 59.9, t _(112)_ = 4.34, p < 0.001). No significant hemispheric differences were observed in the CM-BLA tract (p = 0.11).

#### Robust structural connectivity between the mediodorsal nucleus and the BLA

Similarly, after visually inspecting the medial thalamic nuclei to BLA reconstructions and discarding the unsuccessful ones, we generated MD-BLA streamlines. As mentioned above, the MD comprises the lateral (MDl) and medial (MDm) nuclei together. A robust MD-BLA connection was found, with successful reconstruction in all the participants in the left and in 99% of the participants in the right hemisphere. Notably, a paired sample t-test revealed a significant left hemispheric dominance (t _(112)_ = 3.50, p < 0.001, Cohen’s d = 0.33; see Table 3), consistent with the leftward lateralization observed in other thalamo-amygdala connections.

#### Weak structural connectivity between the inferior colliculus and the BLA

Additionally, we investigated a potentially direct connection between IC and BLA, based on previous evidence in bats (Marsh et al., 2002). The MGN ROI was used as an exclusion mask, to ensure the differentiation of this tract from the previously reconstructed IC-MGN-BLA connection. Our findings revealed connecting streamlines in 82.3% of the participants in the left and in 71.7% of the participants in the right. However, connections among these regions were characterized by a small number of streamlines (Table 3), suggesting weak connectivity. Finally, no significant lateralization was observed in the streamline counts (t _(112)_ = 0.2, p = 0.842, Cohen’s d = 0.02).

### Test-retest reproducibility

To examine the reproducibility of the reconstructed pathways, we used data from 24 participants who underwent a second MRI session, using identical imaging protocol, and evaluated their microstructural (FA correlation coefficients) and macrostructural (Dice index, bundle adjacency, streamline density correlation) results.

Within the posterior thalamus, FA profile correlations exhibited group mean values ranging from 0.73 to 0.98 (Figure 2A) across both hemispheres. Among these connections, the IC-MGN-BLA tract showed bilateral FA correlation values exceeding 0.81, while all PUL-BLA pathways displayed mean values above 0.90, with the exception of the PuA-BLA tract, which did not exceed 0.75. Macrostructural reproducibility indices followed a similar pattern, with group means ranging from 0.49 to 0.90 for Dice index, 0.10 to 1.10 for bundle adjacency and 0.32 to 0.96 for the streamline density correlation (Figure 2B, C, D), respectively. As noted above, macrostructural reproducibility metrics must be interpreted with caution when streamline counts are low, as sparse reconstructions can reduce overlap and spatial correspondence measures. Within this context, our results indicate high and convergent reproducibility for the IC-MGN-BLA, PuM-BLA, PuInf-BLA, and PuL-BLA tracts across all macrostructural indices. Among these, the PuInf-BLA pathway exhibited the highest overall reproducibility, reflected by consistently strong Dice overlap, bundle adjacency, and streamline density correlations. In contrast, the PuA-BLA tract, characterized by a low number of reconstructed streamlines (Table 3), showed the poorest reproducibility across all macrostructural measures. Given the known sensitivity of these metrics to sparse tract reconstructions, these results did not allow definitive conclusions about the reliability of the PuA-BLA connection.

**Figure 2:**
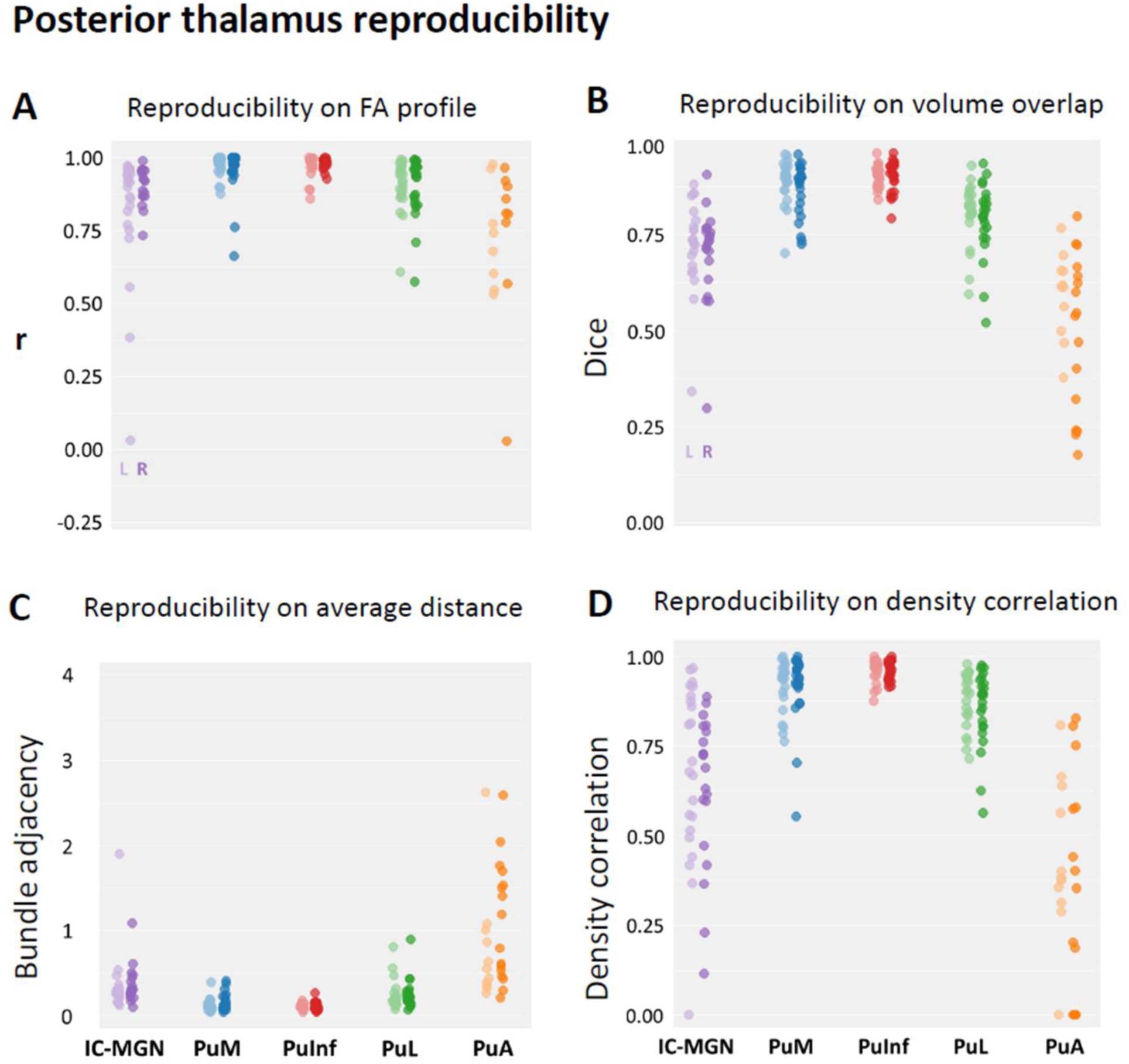
Test-retest reproducibility of tracts connecting the posterior thalamic nuclei to the basolateral amygdala. **(A)** Strip plots showing the distribution of FA correlation coefficients between test and retest for each tract and each subject (lighter color columns represent the left hemisphere; darker color columns represent the right hemisphere). Each dot represents the correlation coefficient for one participant. Agreement indices distribution for the: **(B)** Dice coefficient, **(C)** bundle adjacency and **(D)** density correlation between test and retest for each tract and each subject.

The entire reconstructed IC-MGN-BLA tract of a representative subject is shown in Figure 3A, B, while Figure 3C, D highlights the streamlines passing through the MGN relative to neighboring thalamic nuclei. Figure 4 shows the reliably reconstructed pathways between the PUL and the BLA, illustrating the distinct trajectories of each PUL subregion as they project towards BLA. The pathways arise from distinct subregions of the PUL and converge with slight overlap at their termination points within the BLA. The PuA is excluded from this figure due to its uncertain reconstruction and is instead reported in the Supplemental Figure S1.

**Figure 3:**
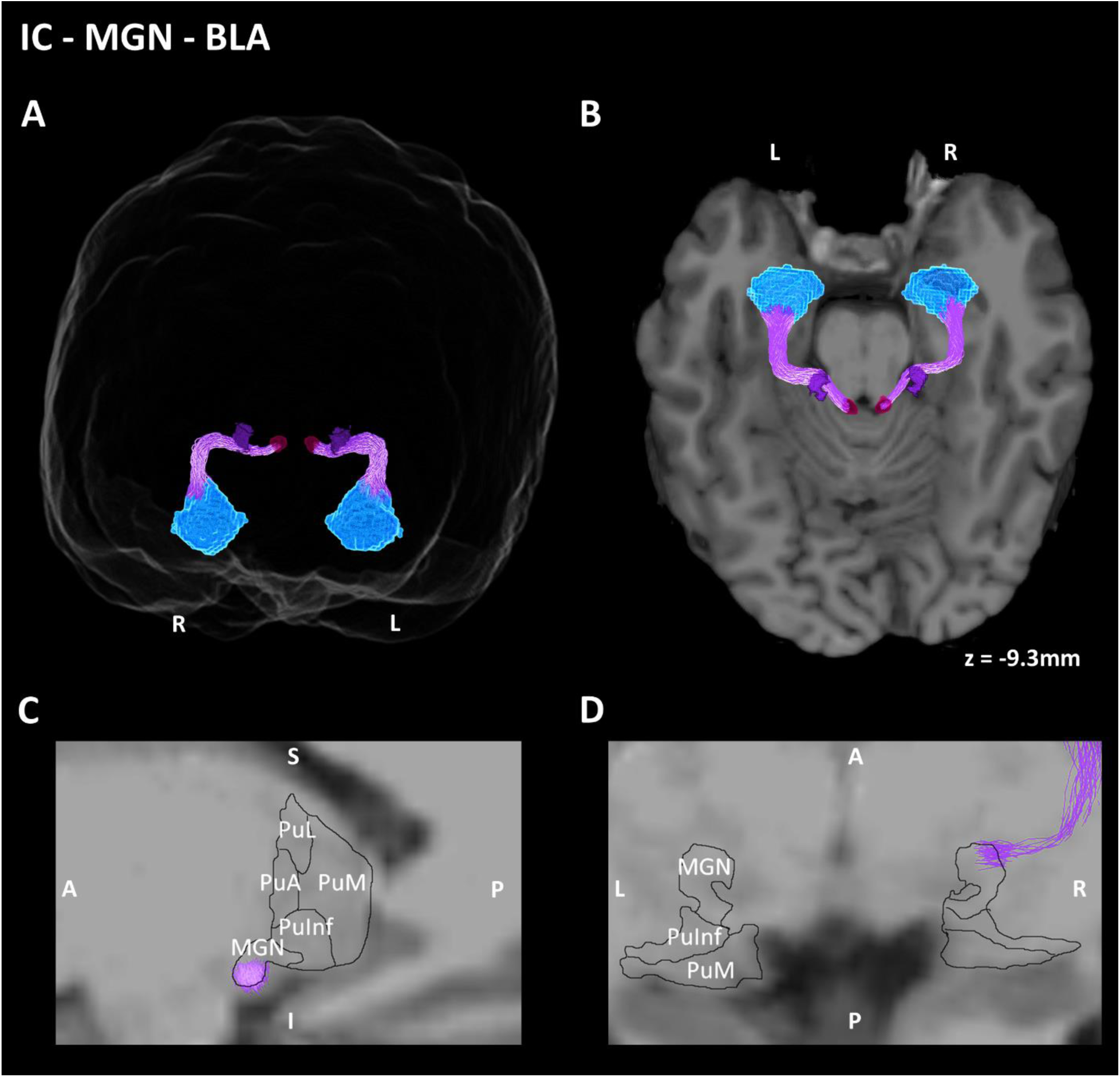
Probabilistic tractography reconstruction of the IC-MGN-BLA pathway in a representative subject. Fibers reconstructed seeding from the inferior colliculus (IC, magenta), passing through the medial geniculate nucleus (MGN; purple) and terminating to the basolateral amygdala (BLA; light blue). 3D representation of the reconstructed pathway in (**A**) a glass brain, (**B**) an axial slice. (**C, D**) Sagittal and axial sections of the thalamus, respectively, are illustrated at higher magnification, showing streamlines passing through MGN.

**Figure 4:**
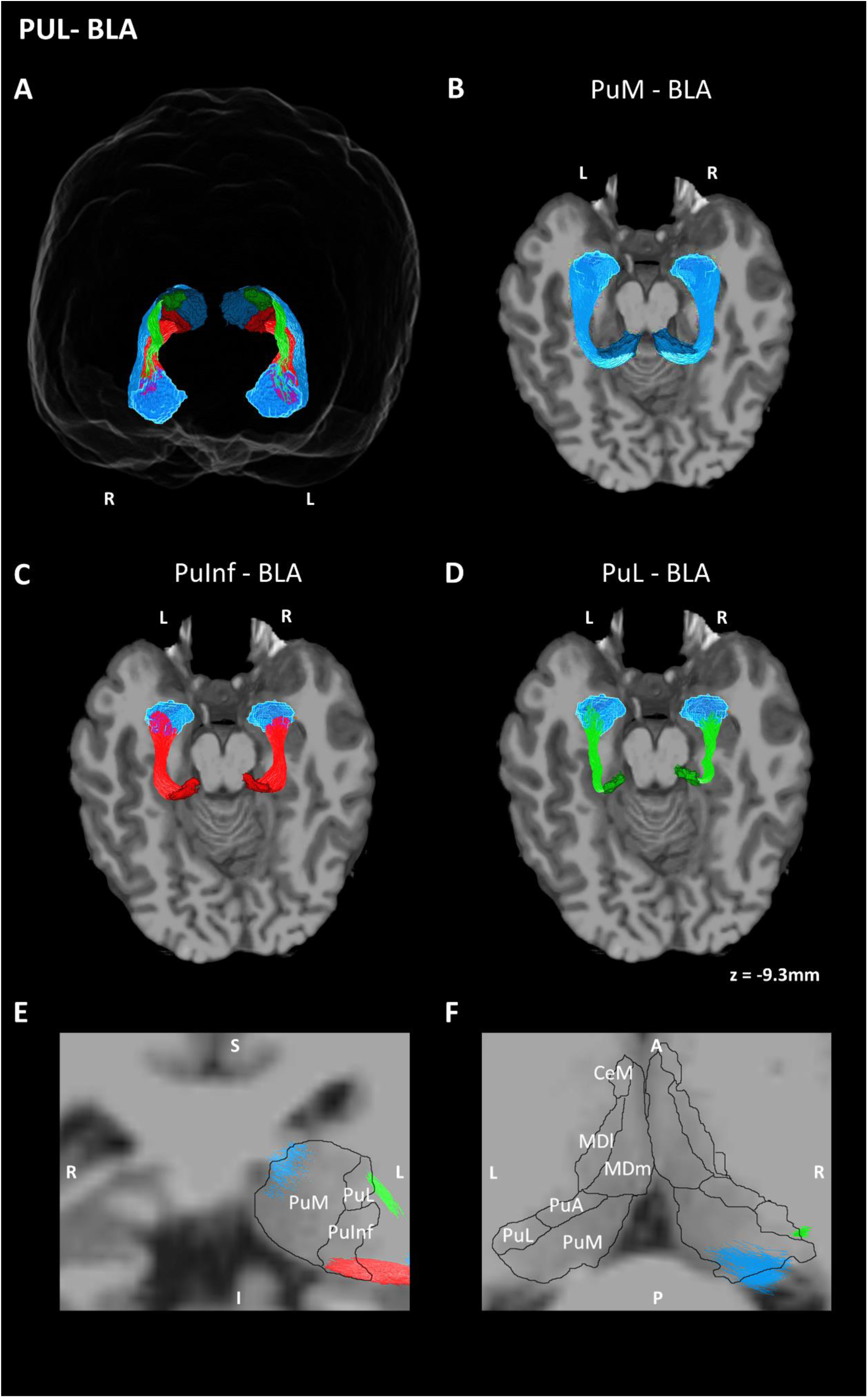
Probabilistic tractography reconstructions of the PUL-BLA tracts in a representative subject. Color scheme is the same as used in Figure 2. Fibers reconstructed seeding separately from the medial (PuM; dark blue), inferior (PuInf; red) and lateral pulvinar (PuL; green) and terminating to the basolateral amygdala (BLA; light blue). (**A**) 3D representation of the reconstructed pathways in a glass brain. 3D representations of the distinct (**B**) PuM-BLA, (**C**) PuInf-BLA and (**D**) PuL-BLA tracts in axial slices. (**E, F**) Coronal and axial sections of the thalamus at higher magnification, showing streamlines originating from distinct PUL subregions.

In turn, intralaminar-BLA projections from the Pf, CeM, and CM nuclei showed moderate microstructural reproducibility (FA correlations 0.60-0.69; Figure 5A) and modest macrostructural indices (Dice 0.44-0.67, bundle adjacency 0.40-0.92, streamline density 0.26-0.54; Figure 5B, C, D). Substantial intrasubject variability resulted in long-tailed distribution patterns (Figure 5). Overall, Pf-BLA and CeM-BLA tracts exhibited moderate reproducibility, while CM-BLA displayed lower but still measurable reproducibility, potentially due to failure of pathway reconstruction, or very little streamlines reconstructed, which limited definitive conclusions about its anatomical validity. Figure 6 illustrates the Pf-BLA and CeM-BLA reconstructions. As it can be observed, both trajectories arise from distinct intralaminar thalamic regions, they partially merge, forming closely adjacent pathways before reaching their termination in the BLA. The CM-BLA reconstructed pathway is shown in Supplemental Figure S1.

**Figure 5:**
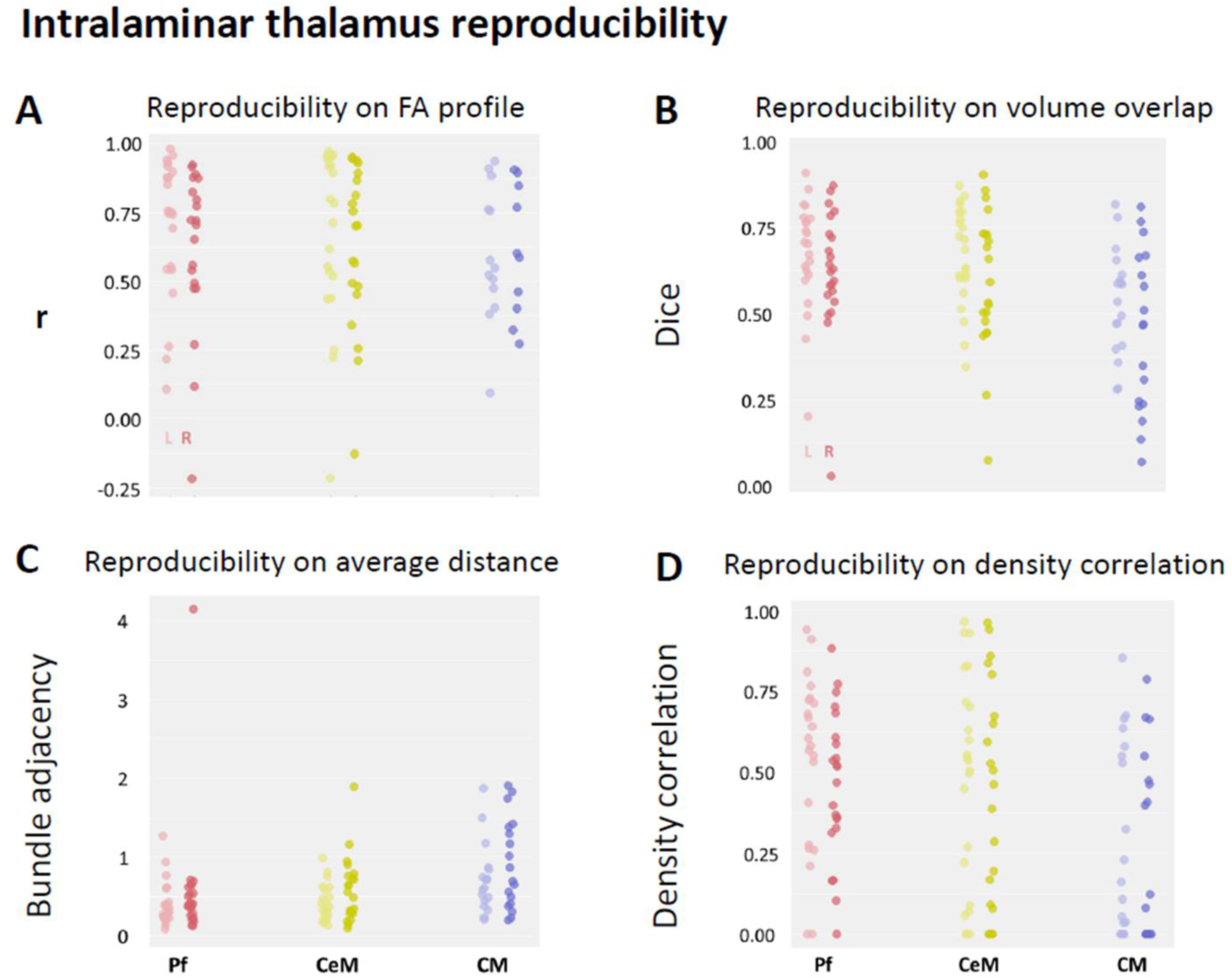
Test-retest reproducibility of tracts connecting the intralaminar thalamic nuclei to the basolateral amygdala. **(A)** Strip plots showing the distribution of FA correlation coefficients between test and retest for each tract and each subject (lighter color columns represent the left hemisphere; darker color columns represent the right hemisphere). Each dot represents the correlation coefficient for one participant. Agreement indices distribution for the: **(B)** dice coefficient, **(C)** bundle adjacency and **(D)** density correlation between test and retest for each tract and each subject.

**Figure 6:**
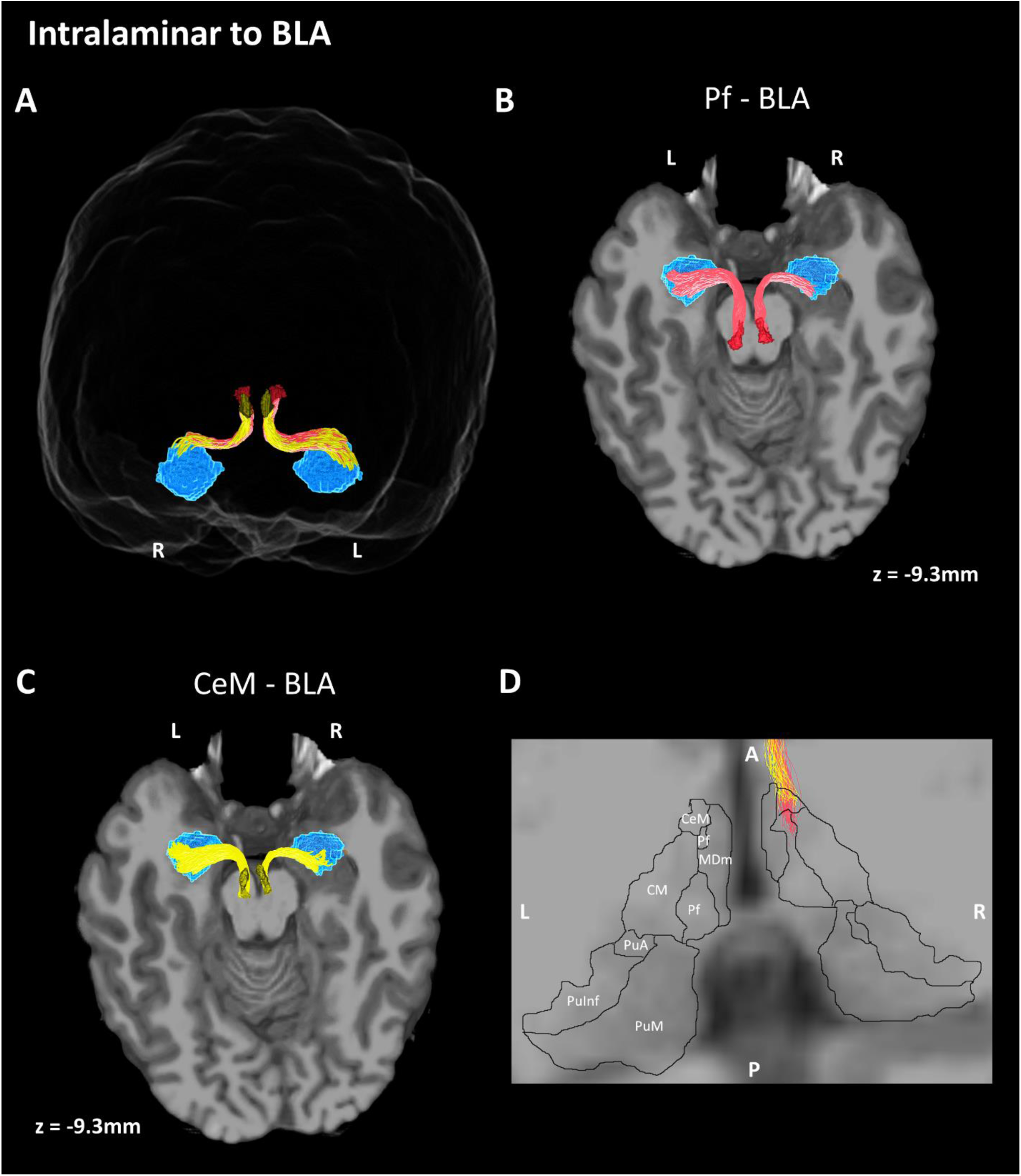
Probabilistic tractography reconstructions of the Pf-BLA and CeM-BLA tracts in a representative subject. Color scheme is the same as used in Figure 5. Fibers reconstructed seeding from parafascicular nucleus (Pf, pink) and centromedial nucleus (CeM; yellow) and terminating to the basolateral amygdala (BLA; light blue). (**A**) 3D representation of the reconstructed pathways in a glass brain. 3D representations of the distinct (**B**) Pf-BLA and (**C**) CeM-BLA tracts in axial slices. (**D**) Axial section of the thalamus at higher magnification, showing streamlines originating from distinct intralaminar subregions.

Meanwhile, the MD-BLA tract exhibited moderate-to-high reproducibility, with mean FA profile correlations of 0.63 (left) and 0.74 (right; Figure 7A). Macrostructural indices were similarly robust (Dice 0.67 for left / 0.65 for right; bundle adjacency 0.49 for left /0.52 for right; streamline density 0.57 for left /0.51 for right; Figure 7B, C, D). These results support a reliable reconstruction of the MD-BLA pathway. Reconstructions (see Figure 8) revealed streamlines originating from both the lateral (MDl) and medial (MDm), notably with the majority arising from the medial magnocellular division (MDm).

**Figure 7:**
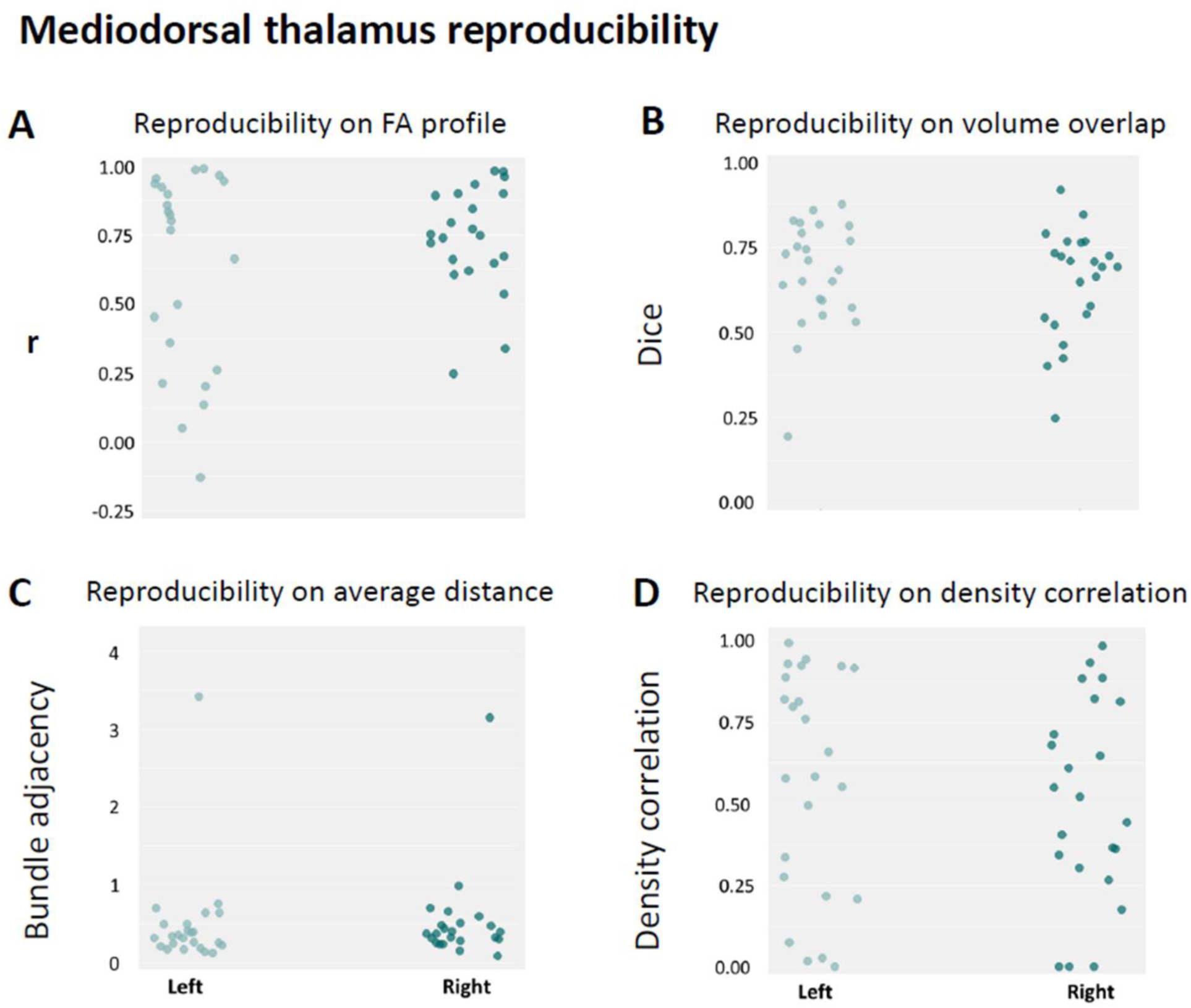
Test-retest reproducibility of tracts connecting the mediodorsal thalamus to the basolateral amygdala. **(A)** Strip plots showing the distribution of FA correlation coefficients between test and retest for each subject (lighter color column represents the left hemisphere; darker color column represents the right hemisphere). Each dot represents the correlation coefficient for one participant. Agreement indices distribution for the: **(B)** Dice coefficient, **(C)** bundle adjacency and **(D)** density correlation between test and retest for each subject.

**Figure 8:**
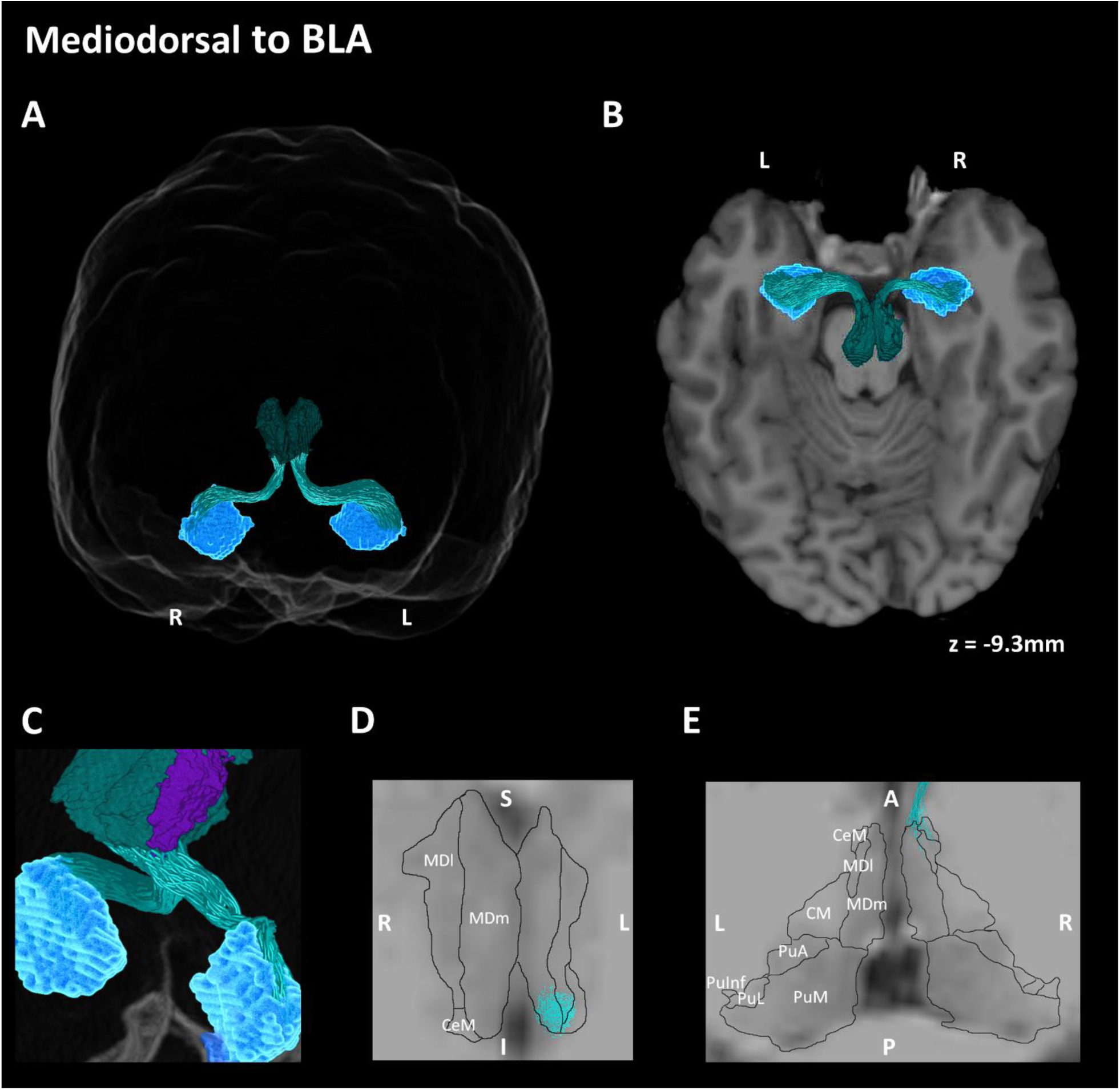
Probabilistic tractography reconstructions of the mediodorsal thalamus to basolateral amygdala tract in a representative subject. Color scheme is the same as used in Figure 7. Fibers reconstructed seeding from mediodorsal nucleus (MD, petrol) and terminating to the basolateral amygdala (BLA; light blue). 3D representation of the reconstructed pathway in (**A**) a glass brain, (**B**) an axial slice. (**C**) 3D visualization illustrating the medial (MDm; petrol) and lateral (MDl; purple) subdivisions of the mediodorsal nucleus in distinct colors, emphasizing their anatomical separation. (**D**, **E**) Coronal and axial sections of the thalamus at higher magnification, showing streamlines originating mainly from the medial magnocellular subdivision (MDm).

Finally, the sparse reconstruction of the IC-BLA pathway in several participants precludes conclusions about the existence of a direct anatomical connection bypassing the MGN in humans. Although FA profile correlations were moderate (0.76 for left /0.77 for right; Supplemental Figure S2A), reproducibility of the right IC-BLA tract could only be assessed in 4 of 24 participants due to insufficient streamlines in both test and retest sessions. Macrostructural reproducibility was consistently low (Dice 0.53 for left /0.47 for right; bundle adjacency 1.04 for left /1.01 for right; streamline density 0.43 for left /0.28 for right; Supplemental Figure S2B, C, D). Given this lack of robust evidence, we include the visual representation of this tract in Supplemental Figure S1. Detailed group mean reproducibility metrics for all tracts are provided in Supplemental Table S1.

## Discussion

We present novel human evidence for the existence of multiple direct thalamo-amygdala connections potentially analogous to the subcortical ‘low roads’ for fear described in non-human species (Aggleton et al., 1980; Doron & LeDoux, 1999; McFadyen et al., 2020; Pessoa & Adolphs, 2010). Using diffusion-weighted imaging data from a 113-participant sample, we implemented an advanced tractography protocol (Lerma-Usabiaga et al., 2023; Liu et al., 2022; Mengxing et al., 2023) to provide an integrated characterization of connections between the posterior, intralaminar and medial thalamic nuclei and the basolateral amygdala (BLA) within a single human cohort. To examine the reliability of the pathways, we evaluated their micro- and macrostructural reproducibility across two scan sessions in a subset of 24 participants. This tractography pipeline, specifically optimized for resolving pathways through regions with complex fiber crossings, enabled the reconstruction of fine-grained thalamo-amygdala connections with unprecedented anatomical detail and reliability.

### Posterior thalamus as a conserved hub for amygdala connectivity

The reconstructed tracts demonstrated highest consistency and reliability for projections from the posterior thalamus to the BLA. Notably, the IC-MGN-BLA tract was present in nearly all participants, highlighting its structural consistency across individuals, and exhibited excellent microstructural reproducibility alongside moderate-to-high macrostructural reproducibility (Figure 2). While previous human diffusion studies have identified MGN-amygdala connections (Keifer et al., 2015; Rafal & Koller, 2025), our results further indicate that the IC-MGN-BLA pathway, a critical component of the rodent aversion system (Bordi & LeDoux, 1994; LeDoux et al., 1984, 1986), may be conserved in humans. This pathway likely functions as a direct auditory route for emotion-related processing, strongly implicated in fear conditioning, consistent with extensive findings across rodents, cats and humans (Keifer et al., 2015; Khalil et al., 2023; LeDoux et al., 1984; Phelps & LeDoux, 2005; Shinonaga et al., 1994).

Still within the posterior thalamus, connections between the medial, inferior and lateral PUL and the BLA were consistently observed across all participants, showing strong and highly reproducible connectivity. In contrast, anterior PUL-BLA tracts were sparse, with low test-retest reproducibility (Figure 2). Overall, a pulvinar amygdala pathway has been widely associated with fear responses and defensive behaviors across species, as well as with rapid and coarse transmission of sensory information to the amygdala during fear, bypassing the sensory cortex (Day-Brown et al., 2010; Doron & LeDoux, 1999; Elorette et al., 2018; Kragel et al., 2021; McFadyen et al., 2019; Rafal et al., 2015; Tamietto et al., 2012; Wei et al., 2015). In addition, this pathway is believed to support residual (presumably non-conscious) fear processing in blind patients with bilateral occipital damage (Tamietto & De Gelder, 2010).

Our results specifically highlight the medial, the inferior and the lateral PUL subdivisions as the primary contributors to pulvinar-amygdala structural connectivity. Although this pattern aligns with prior studies in rodents and other mammals (where the mediodorsal pulvinar has been described as the main source of amygdala projections; Day-Brown et al., 2010; Doron & LeDoux, 1999), as well as with findings from human diffusion imaging (Abivardi & Bach, 2017; McFadyen et al., 2019; Tamietto et al., 2012), most existing studies have examined the pulvinar as a single structure and thus overlook its internal heterogeneity. This distinction is particularly relevant, as pulvinar subdivisions are known to subserve distinct roles in attention and stimulus processing. For instance, while the inferior and lateral PUL may be largely devoted to visual processing, the medial and anterior PUL show involvement in auditory or multisensory integration (see Bourgeois et al., 2020; Froesel et al., 2024). Therefore, the different pulvinar-amygdala connections identified to date may play distinct roles in affective processing, an issue that should be addressed in future research.

### Consistent but variable human intralaminar - BLA projections

Additionally, we explored intralaminar nuclei projections to BLA which have been previously described in rodents, non-human primates and partly in humans (Aggleton et al., 1980; Kumar et al., 2023; Van Der Werf et al., 2002). Unlike the medial thalamic nuclei, the intralaminar group maintains relatively consistent anatomical definitions across species, facilitating cross-species comparisons. Among the intralaminar nuclei examined, the parafascicular (Pf), centromedial (CeM), and centromedian (CM) nuclei displayed the most consistent connectivity with the BLA, although the number of reconstructed streamlines varied across individuals. Notably, Pf and CeM showed moderate-to-high reproducibility, both at microstructural and macrostructurally levels (Figure 5), consistent with their overlapping connectivity profiles attributed to close anatomical proximity in humans (Kumar et al., 2023), as well as prior findings in animal models (Van Der Werf et al., 2002; Vertes et al., 2022). In contrast, the CM-BLA tract exhibited low reproducibility across participants at both levels (Figure 5), potentially reflecting broader projections across amygdala subregions rather than a targeted BLA connection (Vertes et al., 2012). These findings suggest that, similarly as in rodents, the intralaminar nuclei may serve to integrate affective functions, highlighting conserved intralaminar-amygdala connectivity despite thalamic specialization in humans.

### Robust mediodorsal thalamus connections with BLA in humans

We further examined the medial thalamic nuclei based on rodent and non-human primate literature implicating the midline nuclei in fear processing and extinction circuits (Aggleton et al., 1980; Dobi et al., 2013; Van Der Werf et al., 2002). Our tractography revealed robust connectivity between the lateral and medial (magnocellular) MD and the BLA, with streamlines predominantly originating from the medial MD. Studies in rodents support a critical role of MD in fear extinction and the consolidation and retrieval of long-term fear memories (Lee et al., 2011; Li et al., 2004). However, these connections could reflect an amygdala rather than a thalamic output (Abivardi & Bach, 2017), consistent with the strong reciprocal connectivity between MD and prefrontal cortex (Mengxing et al., 2023). As such, MD-BLA connection may mediate output fear responses rather than the processing of fear-related signals (Mátyás et al., 2014). Notably, we detected no significant connections from the paratenial or reuniens nuclei, despite robust evidence for such projections in rodents, likely due to their small size and the resulting tractography limitations (Van Der Werf et al., 2002). Additionally, the connectivity we observed within the medial thalamic ROI may partially reflect contributions from other midline nuclei, such as the paraventricular and intermediodorsal (IMD) nuclei, which are not explicitly delineated in current human atlases but have been shown to project to the BLA in rodents (Dobi et al., 2013; Van Der Werf et al., 2002; Vertes et al., 2022). These limitations underscore the challenge of defining homologous thalamic nuclei across species.

### No reliable IC-BLA direct connections identified in humans

Finally, we tested for a direct IC-BLA pathway, which has been reported in certain non-human species, such as bats (Marsh et al., 2002). This connection was not reliably reconstructed in our dataset, with low or absent streamline counts in the majority of participants. Test-retest macrostructural reproducibility was also low, despite relatively high microstructural reproducibility driven by a small subset of individuals (Figure S1). Previous diffusion MRI studies in tinnitus patients have reported direct IC-amygdala connections using less anatomically constrained tractography approaches (Crippa et al., 2010). Our findings suggest that such direct streamlines may have partially overlapped with an IC–MGN–BLA pathway, as the MGN was not explicitly excluded in their analyses. This interpretation is supported by the robust and consistent reconstruction of the IC-MGN-BLA tract across nearly all participants in our study.

### Leftward hemispheric lateralization in specific tracts

Across thalamic subdivisions, the IC-MGN-BLA, PuInf-BLA, Pf-BLA, CeM-BLA, and MD-BLA tracts were consistently left-lateralized in streamline count. This pattern aligns with prior evidence of left-dominant thalamo-amygdala connectivity (Abivardi & Bach, 2017; Rafal & Koller, 2025), and with lesion and functional studies implicating the left amygdala in processing speech-related threat cues (Frühholz et al., 2015; Pannese et al., 2016). In contrast, leftward lateralization of PUL-BLA pathways is less consistent with previous findings from visual emotion processing (McFadyen et al., 2019). Whether this lateralization reflects pathway-specific functional specialization or broader hemispheric asymmetries in affective processing remains to be determined.

### Conclusions

In sum, we provide a novel, integrated anatomical mapping of multiple thalamo-amygdala pathways in the human brain (Figure 9), revealing direct routes that may be homologous to the subcortical ‘low-road’ circuits extensively described in animal models. These pathways vary in their consistency and reproducibility, with some emerging as highly robust, while others appear weaker or potentially absent. Our findings delineate several conserved thalamo-amygdala connections from posterior, intralaminar and medial thalamic subdivisions, which may support rapid transmission of sensory and affective signals and enable flexible defensive responses rather than redundant processing. Future research is needed to disentangle the specific functional roles of each pathway in human affective processing. Inter-individual variability likely reflects evolutionary specialization of the primate brain, which may render some pathways functionally redundant or absent in certain individuals (McFadyen et al., 2020). Finally, inconsistencies across studies may stem from differences in thalamic parcellation, as nuclei defined in rodents are often subsumed within broader human atlases. Together, this work provides a unified anatomical framework linking animal and human models and supports the existence of conserved, distinct subcortical pathways for affective processing.

**Figure 9:**
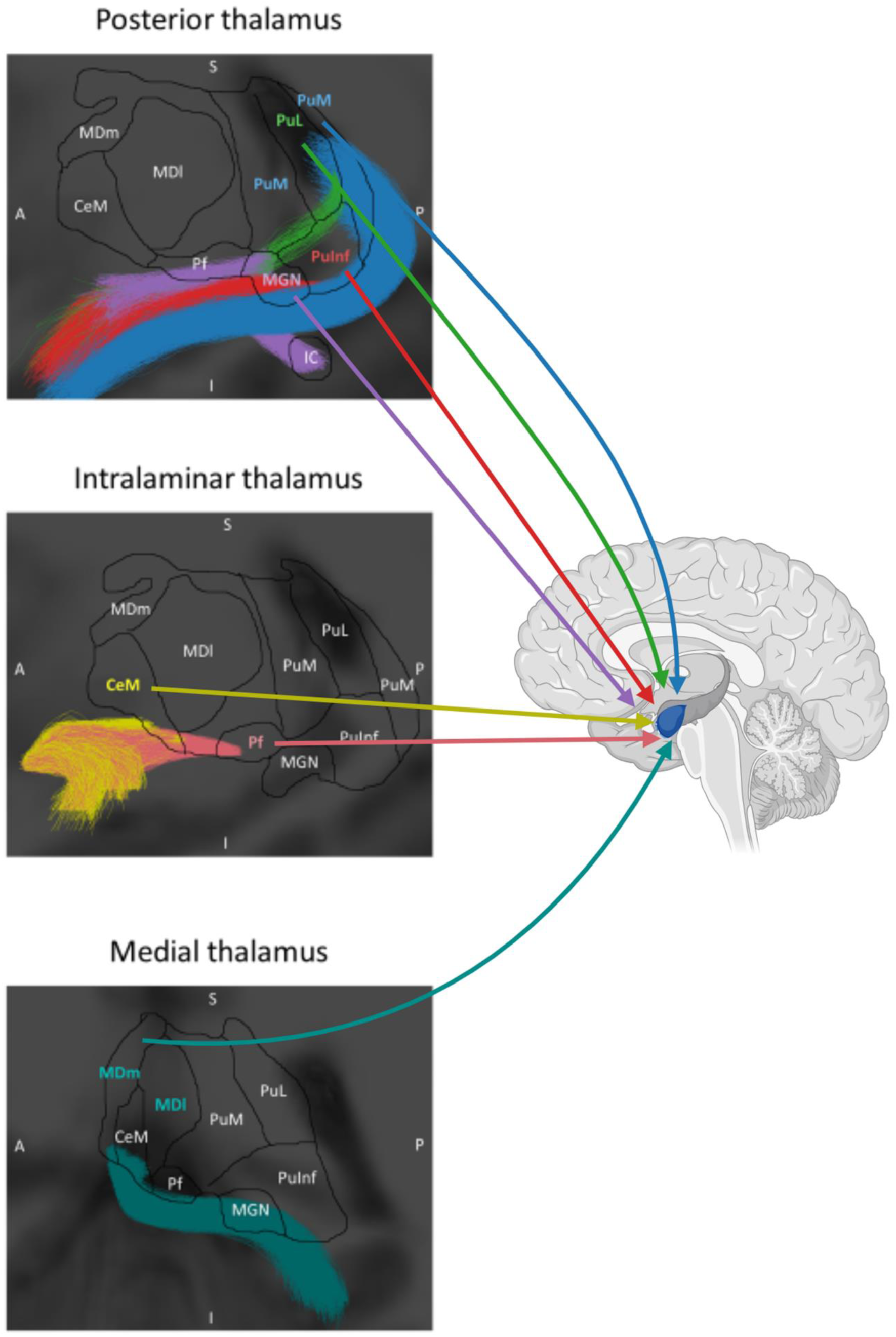
Representation of thalamus-basolateral amygdala (BLA) connections in the human brain. Posterior thalamic projections include streamlines originating from the medial geniculate nucleus (MGN), medial pulvinar (PuM), inferior pulvinar (PuInf), and lateral pulvinar (PuL). Intralaminar thalamic projections include streamlines originating from the parafascicular nucleus (Pf) and central medial nucleus (CeM). Medial thalamic projections include streamlines originating from the mediodorsal medial (MDm) and mediodorsal lateral (MDl) nuclei.

## Acknowledgments

This work was supported by the European Research Council (ERC) under the European Union’s Horizon 2020 research and innovation program (Grant agreement No.101088954), the Spanish Ministry of Science and Innovation (PID2020-116311GA-I00, PRE2021-097083), the María de Maeztu Center of Excellence program (MDM-2017-07-29-20-2), and the Generalitat de Catalunya (SGR2021-00356). P.M.P-A. was supported by a grant from the Spanish Ministry of Science and Innovation (PID2024-159163NB-I00), and acknowledges funding to the BCBL from the Basque Government through the BERC 2022–2025 program and from the Spanish State Research Agency through the Severo Ochoa Center of Excellence accreditation CEX2020-001010-S).

## Supplemental Material

**Figure S1:**
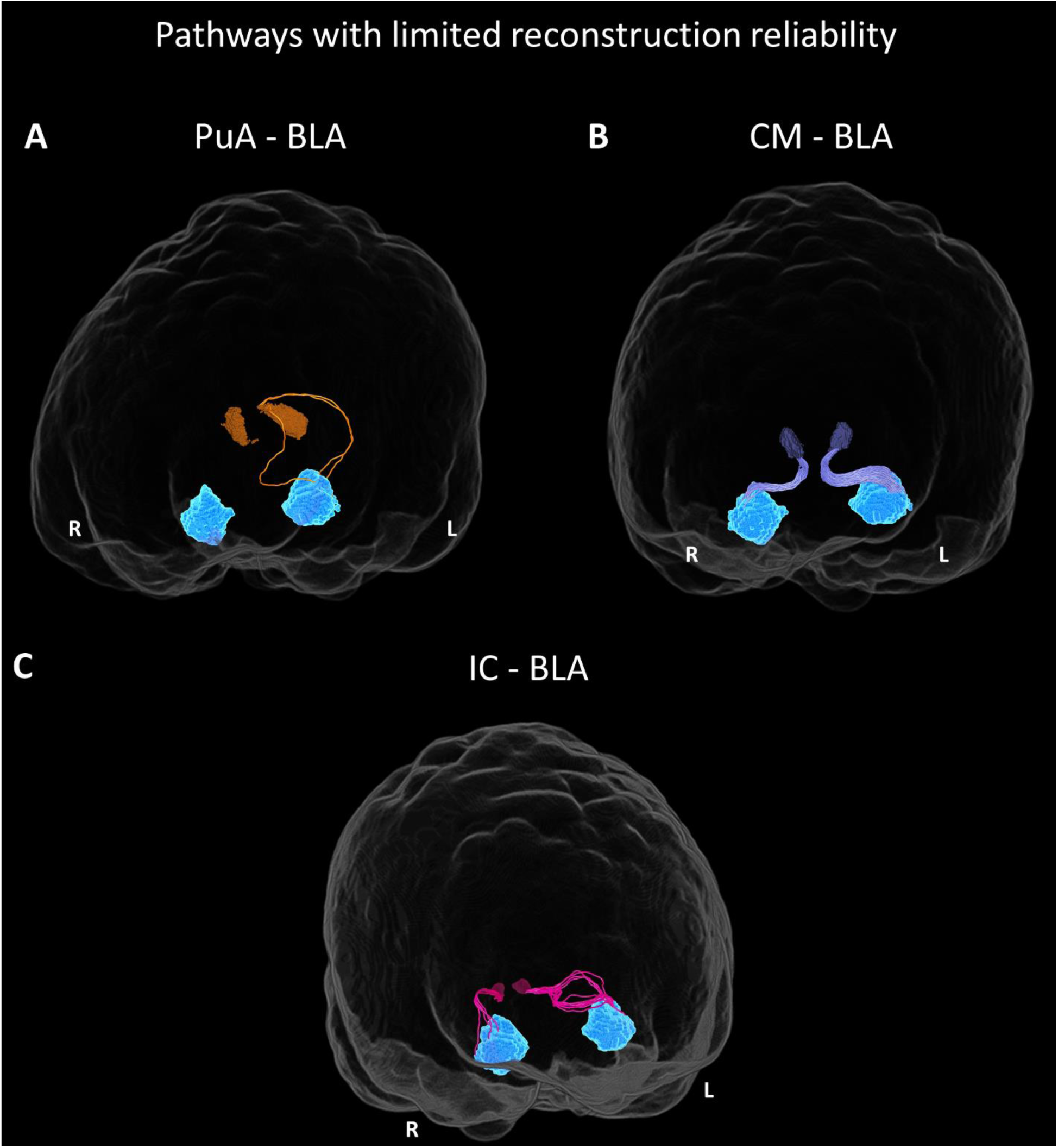
Probabilistic tractography reconstructions of thalamo-amygdala pathways with lower reconstruction consistency in a representative subject. 3D representations show the (A) PuA-BLA pathway seeding from the anterior pulvinar (PuA, orange), (B) CM-BLA pathway seeding from the centromedian intralaminar nucleus (CM, purple), and (C) IC-BLA pathway seeding from the inferior colliculus (IC, magenta) and terminating in the basolateral amygdala (BLA, light blue). These pathways were sparsely reconstructed, reflecting their limited reproducibility across participants.

**Figure S2:**
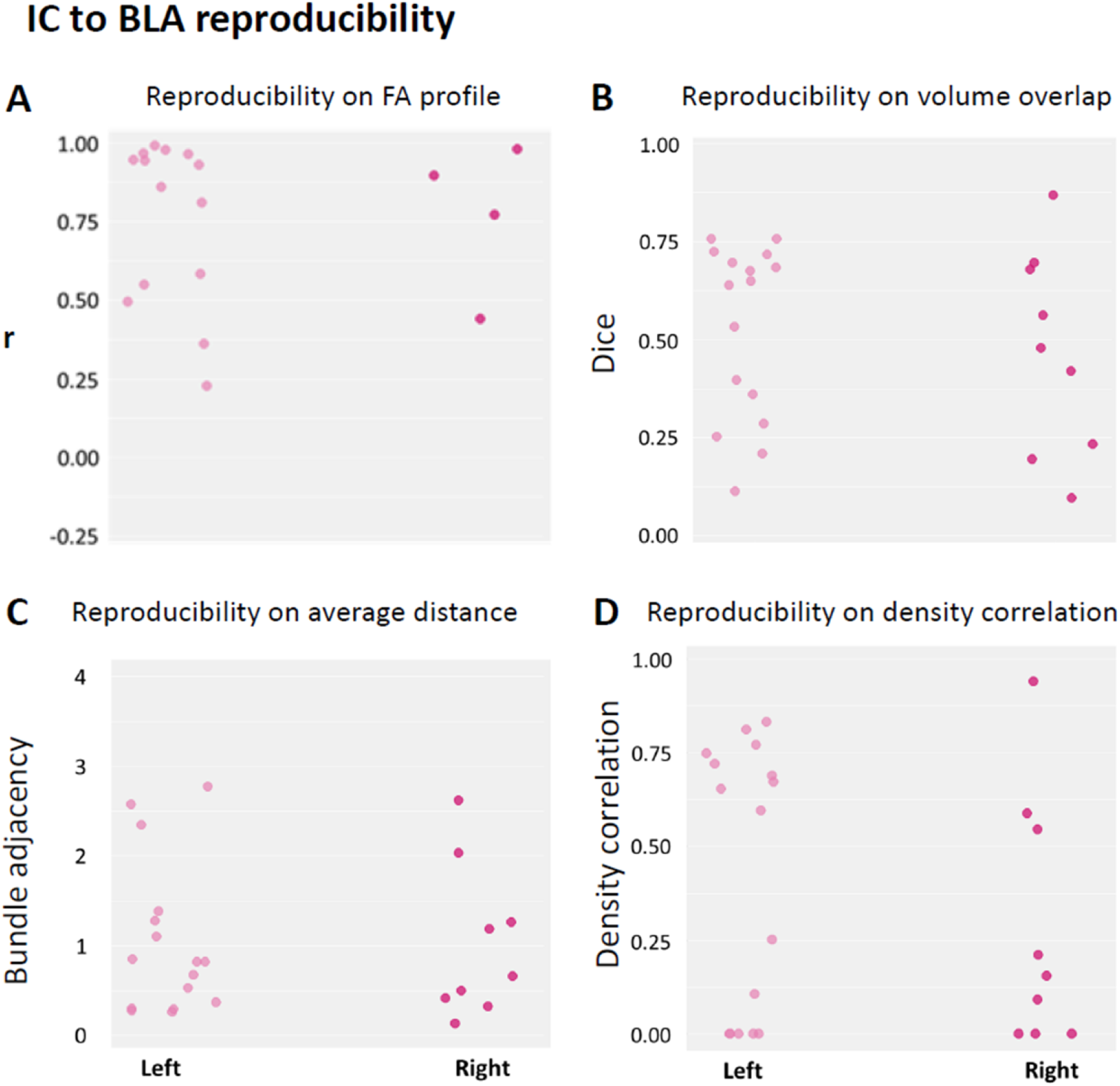
Test-retest reproducibility of the inferior colliculus to basolateral amygdala tract. (**A**) Strip plots showing the distribution of FA correlation coefficients between test and retest for each subject (lighter color column represents the left hemisphere; darker color column represents the right hemisphere). Each dot represents the correlation coefficient for one participant. Agreement indices distribution for the: **(B)** Dice coefficient, **(C)** bundle adjacency and **(D)** density correlation between test and retest for each subject.

**Table S1:**
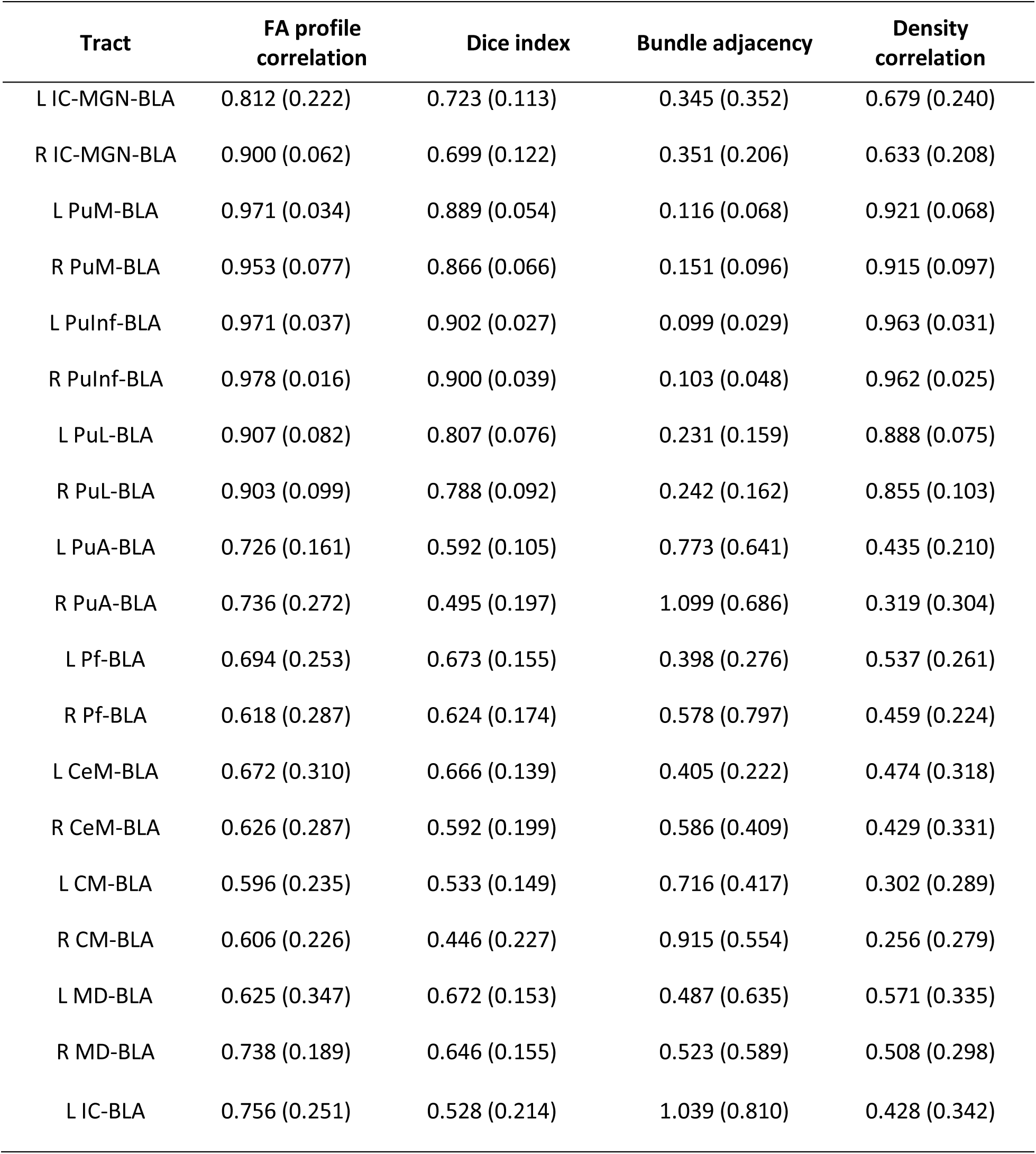
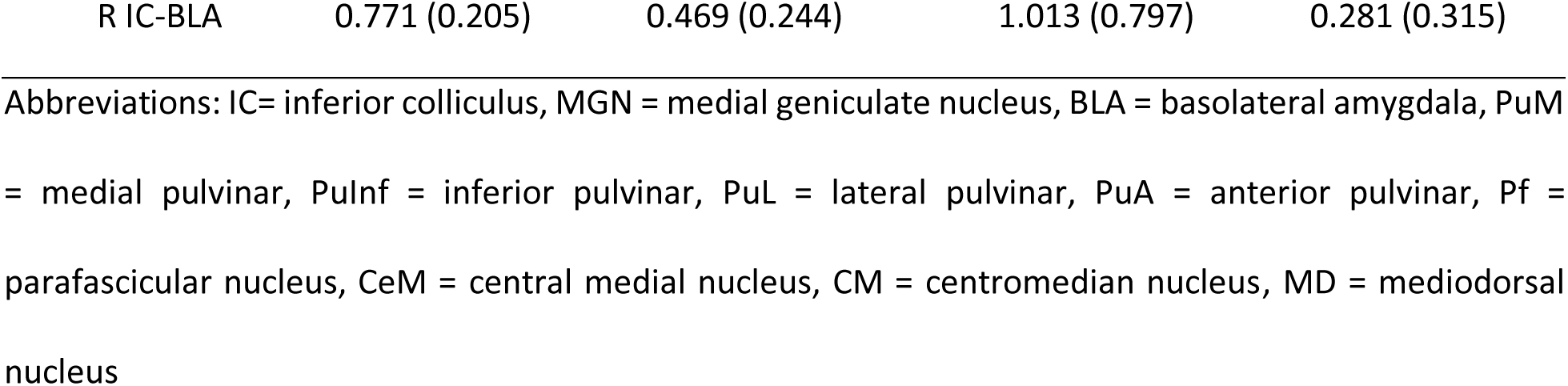
Test-retest reproducibility indices for micro- and macrostructural levels of all tracts bilaterally.

| Tract | FA profile correlation | Dice index | Bundle adjacency | Density correlation |
| --- | --- | --- | --- | --- |
| L IC-MGN-BLA | 0.812 (0.222) | 0.723 (0.113) | 0.345 (0.352) | 0.679 (0.240) |
| R IC-MGN-BLA | 0.900 (0.062) | 0.699 (0.122) | 0.351 (0.206) | 0.633 (0.208) |
| L PuM-BLA | 0.971 (0.034) | 0.889 (0.054) | 0.116 (0.068) | 0.921 (0.068) |
| R PuM-BLA | 0.953 (0.077) | 0.866 (0.066) | 0.151 (0.096) | 0.915 (0.097) |
| L PuInf-BLA | 0.971 (0.037) | 0.902 (0.027) | 0.099 (0.029) | 0.963 (0.031) |
| R PuInf-BLA | 0.978 (0.016) | 0.900 (0.039) | 0.103 (0.048) | 0.962 (0.025) |
| L PuL-BLA | 0.907 (0.082) | 0.807 (0.076) | 0.231 (0.159) | 0.888 (0.075) |
| R PuL-BLA | 0.903 (0.099) | 0.788 (0.092) | 0.242 (0.162) | 0.855 (0.103) |
| L PuA-BLA | 0.726 (0.161) | 0.592 (0.105) | 0.773 (0.641) | 0.435 (0.210) |
| R PuA-BLA | 0.736 (0.272) | 0.495 (0.197) | 1.099 (0.686) | 0.319 (0.304) |
| L Pf-BLA | 0.694 (0.253) | 0.673 (0.155) | 0.398 (0.276) | 0.537 (0.261) |
| R Pf-BLA | 0.618 (0.287) | 0.624 (0.174) | 0.578 (0.797) | 0.459 (0.224) |
| L CeM-BLA | 0.672 (0.310) | 0.666 (0.139) | 0.405 (0.222) | 0.474 (0.318) |
| R CeM-BLA | 0.626 (0.287) | 0.592 (0.199) | 0.586 (0.409) | 0.429 (0.331) |
| L CM-BLA | 0.596 (0.235) | 0.533 (0.149) | 0.716 (0.417) | 0.302 (0.289) |
| R CM-BLA | 0.606 (0.226) | 0.446 (0.227) | 0.915 (0.554) | 0.256 (0.279) |
| L MD-BLA | 0.625 (0.347) | 0.672 (0.153) | 0.487 (0.635) | 0.571 (0.335) |
| R MD-BLA | 0.738 (0.189) | 0.646 (0.155) | 0.523 (0.589) | 0.508 (0.298) |
| L IC-BLA | 0.756 (0.251) | 0.528 (0.214) | 1.039 (0.810) | 0.428 (0.342) |
| R IC-BLA | 0.771 (0.205) | 0.469 (0.244) | 1.013 (0.797) | 0.281 (0.315) |

Abbreviations: IC= inferior colliculus, MGN = medial geniculate nucleus, BLA = basolateral amygdala, PuM = medial pulvinar, PuInf = inferior pulvinar, PuL = lateral pulvinar, PuA = anterior pulvinar, Pf = parafascicular nucleus, CeM = central medial nucleus, CM = centromedian nucleus, MD = mediodorsal nucleus

